# Synapsin-1 and actin form ordered nanoscale assemblies

**DOI:** 10.64898/2026.09.25.754542

**Authors:** Daniel Mansour, Rajdeep Chowdhury, Aravind Chandrasekaran, Tiago Mimoso, Donatus Krah, Aleksandr Korobeinikov, Akshita Chhabra, Dragomir Milovanovic, Sarah Köster, Ali H. Shaib, Silvio O. Rizzoli, Padmini Rangamani

## Abstract

Liquid-liquid phase separation is a vital and ubiquitous principle of subcellular organization. Several actin-binding proteins have recently been shown to undergo liquid-liquid phase separation, forming micron-sized droplets that assemble actin into distinct network shapes. However, our knowledge of how these phase-separated proteins organize actin filaments at the nanoscale is rather limited. Here, we seek to address this knowledge gap by investigating synapsin-1 condensates through a combination of computational simulations and nanoscale imaging experiments. Our results show that while synapsin-1 condensates by themselves lack any special structural organization, the addition of actin filaments results in the generation of a regularly organized actin scaffold within the condensates. The organized scaffold forms both when actin and synapsin-1 are added simultaneously and when actin is added after the formation of synapsin condensates. Dissolving the condensates, by synapsin-1 removal, leaves the actin scaffold largely unaffected in both simulations and experiments, implying that this scaffold should guide droplet reformation when synapsin-1 is added back. Taken together, our findings suggest that the actin-rich component of the condensate retains a structural memory that promotes and guides the reassembly of synapsin-actin condensates. This structural memory may be important in the long-term maintenance of synaptic function *in vivo*.

## Introduction

Liquid-liquid phase separation (LLPS) has emerged as a powerful principle of cellular organization. By concentrating proteins, nucleic acids, and second messengers into dynamic compartments, condensates can create local reaction environments without the need for a surrounding membrane^1–5^. The condensates act as reaction centers, whose organization influences the architecture of other cellular structures, and especially of the cytoskeleton. Several types of condensates have been shown to recruit actin, and to drive its polymerization and bundling^6,7^. Actin filaments group into different shapes ranging from shells to rings, depending on the kinetics of the actin-condensate interactions^8^. Importantly, the condensates do not need to include proteins with a specific polymerase activity, since proteins that only bind actin, without any additional effects, such as lamellipodin, induce actin assembly and bundling^9^. Overall, the main effects induced by the condensates are to concentrate actin, and to provide it with a mechanically deformable interface, which promotes the formation of tightly bundled filament assemblies (e.g., rings^10^). Yet crowding alone cannot account for condensation as proteins such as G3BP2 that non-specifically accumulate actin fail to trigger polymerization^11^. These observations suggest that condensates direct the formation of well-organized actin scaffolds at the mesoscale, mainly at the condensate periphery, at the 1–5 µm scale. However, the nanoscale organization of actin, or of the other proteins, within the fluid condensate phase, remains unknown. In principle, the highly dynamic nature of LLPS-based condensates suggests the possibility that actin filaments within these condensates might be loosely organized. However, no hard evidence for this hypothesis has been generated, mostly due to issues with optical resolution, with condensates appearing to be in homogeneous arrangements when imaged with both conventional imaging optics and super-resolution microscopy tools^12^.

In this study, we seek to understand the nanoscale organization of actin in condensates. As a model protein, we turned to the neuronal protein synapsin-1 because it provides a particularly relevant biological system in which we can understand the nanoscale organization of actin^13^. At the synaptic bouton, synapsins cluster synaptic vesicles, associate with the actin cytoskeleton, and are regulated by phosphorylation during neuronal activity^14–21^. Synapsin-1 undergoes LLPS due to its long and highly positively-charged intrinsically disordered region (IDR) rich in proline and polar residues^22–24^, and also sequesters synaptic vesicles *in vitro*^25–27^. More recently, synapsin-1 was shown to undergo LLPS with synaptic-vesicle-like membranes, supporting the view that synaptic vesicle clusters can be understood, at least in part, as condensate-like assemblies^26,27^. Synapsin-1 has been associated with actin binding for almost four decades^28^, with domain C serving as an actin binding site that is structurally similar to ATP-utilizing enzymes^15,29–31^. Domain C binds both ATP and ADP, with varying affinities, in a calcium-dependent manner^32,33^ and facilitates synapsin-1 multimerization and association with actin and synaptic vesicle phospholipids^14,22–24,34,35^. Synapsin-1 has recently been shown to sequester actin into condensates and induce actin polymerization, forming both rings and aster-like assemblies around synaptic vesicles^11^. However, we still do not understand how actin and synapsin-1 interact at the nanoscale.

In this work, we focused on two main questions: 1.) What is the nanoscale organization of actin filaments within synapsin-1 droplets? and 2.) How do actin structures respond to synapsin-1 droplet dissolution and reformation? To answer these questions and understand the synapsin-actin interaction, we developed a coarse-grained molecular dynamics simulation framework using LAMMPS (Large-scale Atomic/Molecular Massively Parallel Simulator)^36^. LAMMPS provides a highly customizable, particle-based simulation framework useful for coarse-grained molecular dynamics (CGMD) simulations^36^, enabling us to resolve the molecular constituents of the synapsin-1 droplet and its interaction with actin filaments. Simulations from the model predicted that synapsin-1 condensates can organize actin into bundles with high nematic order. Furthermore, we predicted that the actin network can arrange into a well-organized, repetitive scaffold under physiological conditions. To validate this prediction, we turned to wet-lab experiments using reconstituted synapsin-1 and actin. We leveraged an imaging technique called One-step Nanoscale Expansion (ONE) microscopy that provides molecular-scale resolution^37^. ONE microscopy combines the physical expansion of biological samples with a fluorescence fluctuation analysis, to reach lower single-digit nanometer resolutions. This approach enabled us to reveal repetitive actin patterns within the condensates, demonstrating that LLPS-based synapsin-actin assemblies exhibit nanoscale structural organization. In addition, our simulations suggest that, once the condensates form, the actin scaffold is stable and can survive condensate dissolution. This prediction was validated by live imaging approaches. Our findings suggest that synapsin-1 may initially drive the formation of actin scaffolds in synapses, and the same scaffolds later provide a form of local (bouton-specific) memory for the maintenance of synaptic function, throughout activity and plasticity.

## Results

We constructed a physics-based model representing proteins as Lennard-Jones (LJ) particles^38^ to understand how synapsin-synapsin and synapsin-actin interactions drive bundling and condensate formation. Experiments have shown that synapsin condensates are on the order of several hundred nanometers (∼100-500 nm), which these particle-based approaches describe well. Here, we employ a mesoscopic model that represents synapsin, actin, and PEG with spherical LJ particles using LAMMPS (**Fig. 1A**), a widely used molecular dynamics program^39^. Briefly, we modeled synapsin and PEG as spherical particles, and introduced actin as preformed filaments (a string of spherical particles with bonded interactions) with stretching and bending potentials chosen to maintain short, stiff filaments. For simplicity, we assumed that preformed actin filaments are sequestered into synapsin droplets, and we did not model actin polymerization nor filament nucleation. The specific details of the model setup and its parameters are specified in **Supplementary Table 1**. The LAMMPS simulations were performed on the Triton Shared Computing Cluster (TSCC) at the San Diego Supercomputer Center (SDSC)^40^. First, we developed our computational model of a synapsin condensate using a system that consists of synapsin and PEG particles, while modulating the synapsin volume fraction and the synapsin-synapsin interaction strength (ε_Syn-Syn_) (**Fig. 1B**). We used the radius of gyration of the synapsin particles, normalized to the maximum radius measured across all simulation conditions, to measure the degree of coalescence, where smaller values indicate a tighter spherical droplet. This analysis shows that synapsin particles are diffuse at low ε_Syn-Syn_ and begin to coalesce as ε_Syn-Syn_ increases (**Fig. 1C**). Our simulations show that increasing the synapsin volume fraction and increasing ε_Syn-Syn_ lead to the formation of robust droplets in the presence of PEG. Beyond ε_Syn-Syn_ = 2.0, increases in ε_Syn-Syn_ do not result in significant changes to synapsin droplet formation. Thus, we established the regime in which the transition from diffuse to aggregated synapsin occurs within our simulation framework while demonstrating that our physics-based model was capable of capturing the coalescence of synapsin particles to form droplet-like aggregates, given a sufficiently large ε_Syn-Syn_. The droplets formed in our simulations are visually similar to the condensates formed in experiments with synapsin-PEG mixtures (**Fig. 1D, Supplementary Fig. 1**).

**Figure 1:**
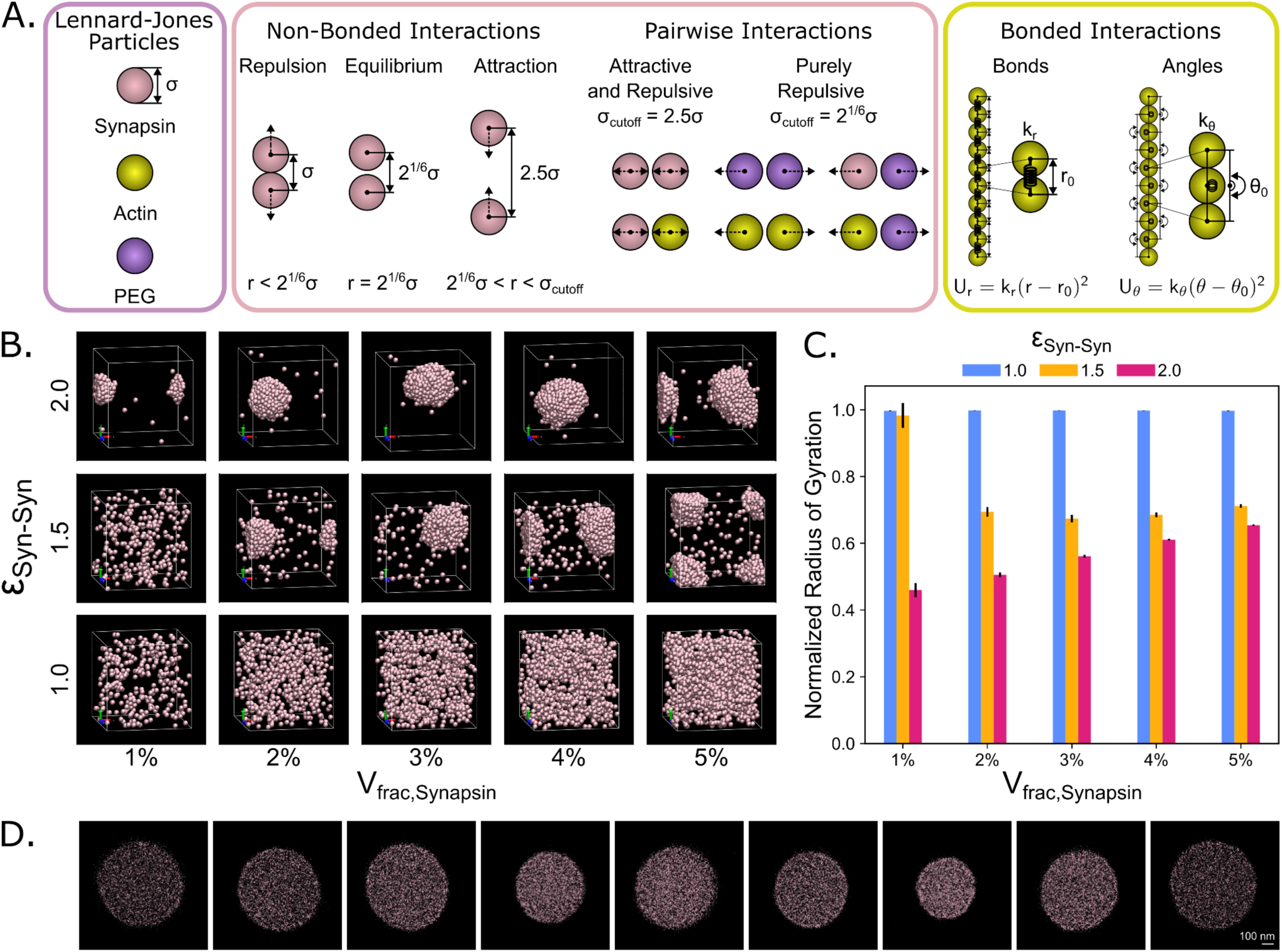
Increasing the strength of Synapsin-Synapsin interactions promotes droplet formation and the strength of coalescence. **A)** Schematic explaining the LAMMPS model setup. Synapsin, actin, and PEG are represented as LJ particles (purple box). Note that synapsin and actin are modeled with the same diameter (σ_Syn-Syn_ = σ_Act-Act_ = 1.0σ) while PEG is modeled with a smaller diameter (σ_PEG-PEG_ ≈ 0.8σ). The pairwise non-bonded interactions (pink box) between particles are governed by the Lennard-Jones equation. The choice of cutoff radius σ_cutoff_ allows us to truncate the range of the LJ potential such that specific pairwise interactions are purely repulsive (σ_cutoff_ = 2^1/6^σ) or both attractive and repulsive (σ_cutoff_ = 2.5σ). Here, synapsin-synapsin and synapsin-actin interactions are both attractive and repulsive while all other interaction pairs are purely repulsive. Actin filaments are composed of a series of actin beads governed by bonded interactions (yellow box) that use harmonic potentials to enforce an equilibrium bond distance and angle between beads within the same actin filament. Please refer to **Supplementary Table 1** and the Methods for a detailed list of parameters used and specific details of the model setup. **B)** Representative final snapshots of simulations with synapsin and PEG particles. The strength of Synapsin-Synapsin interactions is varied by row, while the volume fraction of synapsin is varied by column. The volume fraction of PEG is kept constant at 3%. PEG is not visualized in these snapshots. **C)** Bar chart depicting the mean normalized radius of gyration shows the coalescence of Synapsin particles as the strength of the Synapsin-Synapsin interaction is increased. The error bars represent the standard deviation. *n* = 10 replicates, data from 10 time points in the last 100 frames of the simulations. **D)** Representative image gallery of condensates consisting of synapsin and PEG (the latter is not labeled). Scale bar = 100 nm. Please see **Supplementary Figure 1** for the extended gallery of synapsin condensate images.

### Actin-synapsin composites result in the emergence of nematic order

We next asked how preformed actin filaments interact with the synapsin droplet. To answer this question, we set up simulations in which we fixed the effective synapsin-synapsin interaction strength at ε_Syn-Syn_ = 2.0 to capture the weak multivalent interactions that drive synapsin LLPS, and set the synapsin-actin interaction strength to ε_Syn-Actin_ = 5.0 to ensure strong actin filament recruitment and crosslinking within the synapsin droplet environment. We first simulated the simultaneous addition of actin with synapsin and PEG (**Fig. 2A**) and found that actin and synapsin assemble into rod-like bundles and branched networks that span the periodic boundaries (**Fig. 2A**). We also simulated the case where actin filaments are added after synapsin droplets are formed (**Fig. 2B**). Each simulation condition was replicated 10 times. While the structures from delayed actin addition also formed bundle-like structures (**Fig. 2B**), we identified specific changes to actin filament organization which we will outline below. To quantify the organization of these actin bundles, we calculated the nematic order (S) of all actin filaments in the system, a metric that reports the degree of orientational alignment of anisotropic particles (**Fig. 2C**). Briefly, a low nematic order, roughly S < 0.3, indicates that the filaments are in a disordered phase while a high nematic order, roughly S > 0.8, indicates that the filaments are in an ordered phase. Please note that the disordered phase is characterized by the random orientation of anisotropic actin filaments and is conceptually different from the intrinsically disordered regions of the synapsin protein that are characterized by the lack of a stable 3D protein structure. In between the ordered and disordered phases, roughly 0.3 < S < 0.8, is a partially ordered phase where filaments have an intermediate level of orientational organization and maintain a common preferred direction. We observed that the actin-synapsin composite environment drives the emergence of nematic order in actin filaments across all the simulation conditions tested (**Fig. 2D-F**, **Supplementary Fig. 2**).

**Figure 2:**
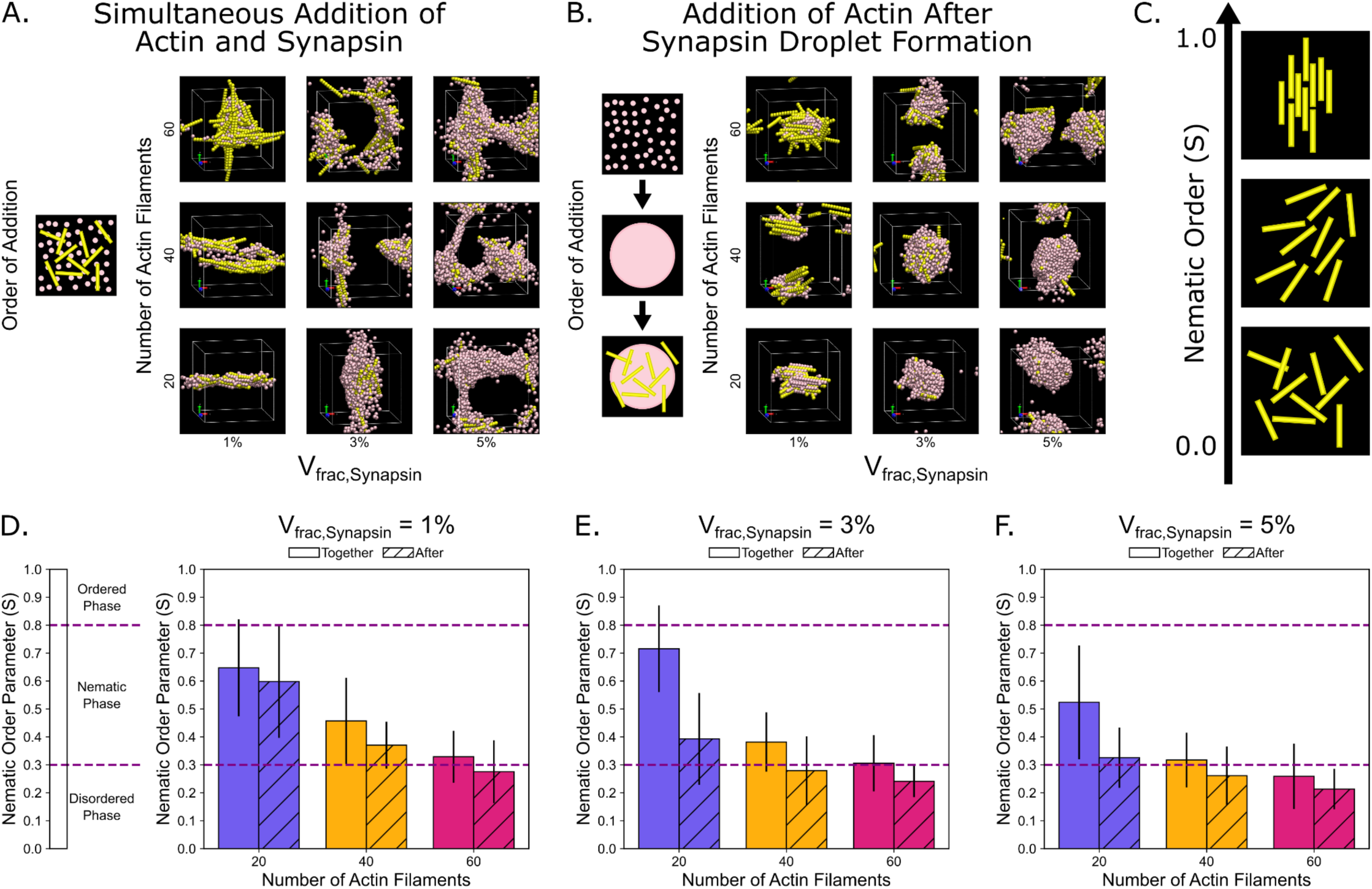
Actin-Synapsin composites result in the emergence of nematic order. **A)** Representative final snapshots for simulation conditions with the simultaneous addition of actin and synapsin. The number of actin filaments in the simulation is varied by row, while the volume fraction of synapsin is varied by column. The volume fraction of PEG is kept constant at 3%. PEG is not visualized in these snapshots. **B)** Representative final snapshots for conditions where actin is added to the system after synapsin droplet formation. The number of actin filaments in the simulation is varied by row, while the volume fraction of synapsin is varied by column. The volume fraction of PEG is kept constant at 3%. PEG is not visualized in these snapshots. **C)** Cartoon of actin filaments ranked by nematic order parameter. **D-F)** Bar chart depicting the mean nematic order parameter and comparing between simulations where actin filaments are added together with synapsin (empty bars) and after synapsin (diagonal stripes), for a synapsin volume fraction of **D)** 1%, **E)** 3%, and **F)** 5%. The error bars represent the standard deviation. *n* = 10 replicates, data from 10 time points in the last 100 frames of the simulations. Please see **Supplementary Figure 2** for the full range of conditions and **Supplementary Figure 3** and **Supplementary Table 2** for detailed statistical analysis.

To understand the relationship between the nematic order and the concentrations of actin and synapsin, we also varied the number of actin filaments and the synapsin volume fraction (**Fig. 2D-F, Supplementary Fig. 2**). Based on Onsager hard-rod theory^41^, we would expect that the nematic order should increase with the number of actin filaments^42^. Surprisingly, we find that this is not the case. Instead, we found that the average nematic order decreased as the number of actin filaments was increased (**Fig. 2D-F**, **Supplementary Fig. 3**). Note that **Figure 2D-F** are select comparisons for the purposes of showing nematic order. Please see **Supplementary Figure 2** for the full range of conditions and **Supplementary Figure 3** and **Supplementary Table 2** for detailed statistical analysis. Closer observation of the actin structures emerging from our simulations reveal that synapsin-mediated crosslinking of actin at high densities caused actin filaments to form more intersections with each other that could become kinetically jammed (**Supplementary Fig. 4**). This jamming locked the filaments into a disordered network state and prevented the emergence of nematic alignment of actin filaments in the system. When we analyzed the results from simulations that had actin filaments added after the synapsin droplet coalesced, we found that the actin structures consistently had a lower average nematic order than the equivalent conditions with simultaneous actin and synapsin addition (**Fig. 2D-F**, **Supplementary Fig. 2**). This lowered nematic order is consistent with the kinetic jamming of actin filaments. Since synapsin is attractive to both itself and actin, synapsin functions as a glue that drives filament organization through crosslinking. When actin and synapsin are added to the system simultaneously, the initially random and diffuse distribution of synapsin provides actin filaments with the freedom to align along extended network branches without strong spatial confinement. Synapsin then coats and coalesces around the actin filaments to guide and stabilize them into rod-like bundles and branched structures. In contrast, introducing actin to a pre-coalesced synapsin droplet forces the filaments into a dense and attractive synapsin environment. The stronger synapsin-actin interaction causes synapsin to readily coat the actin filaments as they enter the droplet, which kinetically traps the filaments in arbitrary orientations and creates a large enough energetic barrier to oppose rotational realignment into a higher nematic order configuration. Thus, the combination of synapsin-actin and synapsin-synapsin interactions in the system give rise to the emergence of a wide range of parameter spaces for the nematic alignment of actin filaments.

We additionally varied the interaction energy between synapsin-actin and synapsin-synapsin to investigate how these parameters would alter the emergence of nematic order. We modulated the ε_Syn-Syn_ and ε_Syn-Actin_ interaction parameters, and measured the resulting nematic order of the actin filaments (**Supplementary Fig. 5**). We selected an intermediate synapsin volume fraction (3%) and chose two cases for a lower (15) and higher (40) number of actin filaments. As before, we also varied the order of actin addition and the number of actin filaments. We show these results as heatmaps that showcase the variability in nematic order across the range of interaction parameters (**Supplementary Fig. 5**). This variability is observed in our simulations through the wide range of actin architectures that emerged within synapsin droplets that depended on parameters such as the actin-synapsin ratio, actin concentration, and relative interaction strengths (**Supplementary Fig. 6**). Despite this large variability, we observed that each heatmap contains a region that supports a higher nematic order (about S ≥ 0.8 for **Supplementary Fig. 5A-C** and S ≥ 0.6 for **Supplementary Fig. 5D**). This region is larger when actin is added simultaneously with synapsin (**Supplementary Fig. 5A-B**) than when actin is added after synapsin droplet formation (**Supplementary Fig. 5C-D**). Additionally, nematic order tended to be higher in systems with fewer (15) filaments (**Supplementary Fig. 5A,C**) than systems with more (40) filaments (**Supplementary Fig. 5B,D**), which is consistent with the results from **Figure 2**. The region of elevated nematic order was also limited to conditions where ε_Syn-Act_ ≥ ε_Syn-Syn_ and for sufficiently large ε_Syn-Act_, appearing above ε_Syn-Act_ = 1.0 when actin was added simultaneously with synapsin.

### Actin-synapsin composites result in the emergence of regularly organized structures

In many simulation conditions in **Figure 2A**, we noticed that actin formed architectures that collectively spanned across the periodic boundaries, as opposed to actin structures that could be entirely contained within the size of the simulation box frequently seen in the simulation conditions in **Figure 2B**. These structures appeared to have different visual characteristics and various nematic orders. As such, we wondered whether the actin structures with different nematic orders represent different higher-order structures. We compared structures formed in two extreme conditions: actin bundles with high nematic order (N_actin_ = 20, V_frac,Synapsin_ = 1%) (**Fig. 3A**, **Supplementary Fig. 7A**) and actin structures with low nematic order (N_actin_ = 60, V_frac,Synapsin_ = 5%) (**Fig. 3B**, **Supplementary Fig. 7B**). To better visualize the higher-order structures, we took advantage of the periodic boundary conditions to draw the periodic images once in each direction of the original cell to produce a 3×3×3 visualization of the larger structure (**Fig. 3A-B**, **Supplementary Fig. 7A-B**). We found that for the high nematic order condition (N_actin_ = 20, V_frac,Synapsin_ = 1%), the periodic image revealed the formation of a one-dimensional rod-like structure (**Fig. 3A**). Out of the 10 replicates, 6 replicates formed similar one-dimensional rod-like structures and 4 replicates did not form any discernible higher-order structure (**Fig. 3A, Supplementary Fig. 7A,C**). When we repeated the same analysis with the low nematic order condition (N_actin_ = 60, V_frac,Synapsin_ = 5%), we observed that regularly organized structures emerged in all replicates, where 6 replicates formed three-dimensional connected structures while 4 replicates formed two-dimensional sheet-like structures (**Fig. 3B**, **Supplementary Fig. 7B-C**). To ensure the resulting structures formed stably, we used the final structure from the existing simulation as the initial condition of the new simulation, where the original topology is stitched together and arranged into a 2×2×2 simulation box. In the high nematic order condition (N_actin_ = 20, V_frac,Synapsin_ = 1%), the rod-like structures underwent some minor rotation along their axes but remained largely consistent, as shown in the simulation visualization of actin and the 2D maximum intensity projection combined from both actin and synapsin (**Fig. 3C-D**). In the low nematic order condition (N_actin_ = 60, V_frac,Synapsin_ = 5%), the 3D structure remained largely stable, as shown in the simulation visualization of actin and the 2D maximum intensity projection combined from both actin and synapsin (**Fig. 3E-F**). From the 2D maximum intensity projections, we drew a line scan across the evident features and plotted the 1D spatial autocorrelation along the line scan to characterize the repeating spatial organization of actin and synapsin individually for the high nematic order condition (N_actin_ = 20, V_frac,Synapsin_ = 1%) (**Fig. 3G-H**) and the low nematic order condition (N_actin_ = 60, V_frac,Synapsin_ = 5%) (**Fig. 3I-J**). The line scans and autocorrelation show that in both conditions, actin and synapsin localize near each other and that the stitched simulations maintained a regularly organized pattern. Thus, our simulations predict that conditions that give rise to intermediate nematic order of actin filaments are favorable for the emergence of higher order synapsin-actin regularly organized structures.

**Figure 3:**
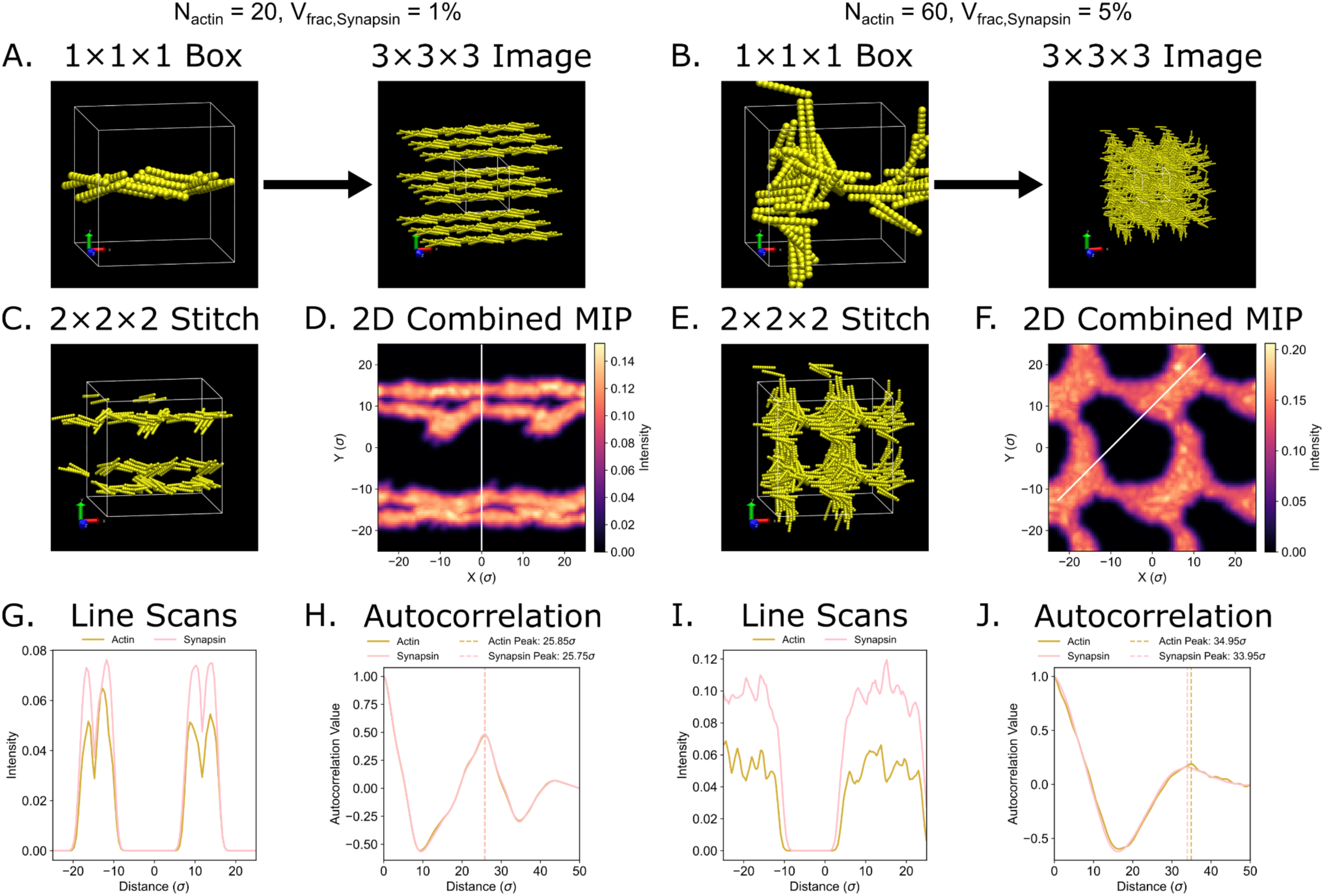
Actin bundles form higher order networks in simulations. **A-B)** Representative final snapshots depicting the actin structure formed in simulations with simultaneous actin and synapsin addition for conditions with **A)** N_actin_ = 20 and V_frac,Synapsin_ = 1%, and **B)** N_actin_ = 60 and V_frac,Synapsin_ = 5%. The volume fraction of PEG is kept constant at 3%. Synapsin and PEG are not visualized in these snapshots. Periodic images are created by a 3×3×3 periodic extension of the simulation boxes to visualize the regularly organized structure formed by actin. **C)** Final snapshot of a 2×2×2 simulation stitched together from the final condition from panel **A** (N_actin_ = 20, V_frac,Synapsin_ = 1%) and run independently to show the stability of the regularly organized structure. **D)** 2D maximum intensity projection along the X-Y plane for combined actin and synapsin signals from the 2×2×2 stitched simulation in panel **C**. **E)** Final snapshot of a 2×2×2 simulation stitched together from the final condition from panel **B** (N_actin_ = 60, V_frac,Synapsin_ = 5%) and run independently to show the stability of the regularly organized structure. **F)** 2D maximum intensity projection along the X-Y plane for combined actin and synapsin signals from the 2×2×2 stitched simulation in panel **E**. **G)** Line scan showing the intensity of the actin (yellow) and synapsin (pink) signals along the indicated region of interest. The white line in panel **D** indicates the line scan drawn across evident features. **H)** 1D spatial autocorrelation on the line scan region in panel **G** detects a regular pattern for both actin (25.85σ) and synapsin (25.75σ), as indicated by the vertical dashed line. **I)** Line scan showing the intensity of the actin (yellow) and synapsin (pink) signals along the indicated region of interest. The white line in panel **F** indicates the line scan drawn across evident features. **J)** 1D spatial autocorrelation on the line scan region in panel **I** detects a regular pattern for both actin (34.95σ) and synapsin (33.95σ), as indicated by the vertical dashed line.

### Actin-synapsin composites show evidence of periodic structure formation *in vitro*

We next asked whether the higher-order actin organization observed in the simulations could also be detected in wet-lab experiments, *in vitro*. To investigate the nanoscale structure of actin-synapsin condensates, we turned to expansion microscopy, in which the specimens are embedded in a swellable gel and are physically expanded, before imaging^43^. We relied on a gel composition that enables a 10-fold expansion^44^, leading to a corresponding increase in imaging resolution, to approximately 25 nm. To bring the overall imaging precision to the molecular level, we combined the 10-fold expansion with the one-step nanoscale (ONE) microscopy procedure, in which the expanded samples are imaged and processed using a fluctuation analysis^37^. ONE microscopy reaches a lateral resolution of ∼1 nm, enabling us to probe the dense space of the condensates.

We first analyzed condensates consisting of synapsin-1 alone (**Fig. 4A**), which we employed to optimize the expansion procedure (**Supplementary Fig. 8**). When analyzed by ONE microscopy, synapsin-1 did not appear to form any special arrangements, with molecules arranged randomly throughout the condensates (**Fig. 4A,D**; see also the image gallery in **Supplementary Fig. 1**). We then combined synapsin-1 and actin, aiming to reproduce, as close as possible, the conditions from the computational simulations presented above. To visualize actin optimally, we relied on a biotinylated version, which was revealed by fluorescently-conjugated streptavidin, before expansion. To mimic the simultaneous addition of actin filaments and synapsin, we incubated synapsin-1 condensates with an actin population that was pre-incubated for 30 minutes at room temperature, to enable it to form filaments in an accelerated exponential fashion, immediately upon entering the condensates. To mimic the delayed incubation of actin filaments with condensates, we used an actin population that was incubated on ice, favoring a prolonged lag phase between actin entering the condensates and the formation of the filaments. At a mesoscopic level, both of these treatments resulted in circular condensates, containing both synapsin-1 and actin (**Supplementary Fig. 9**).

**Figure 4:**
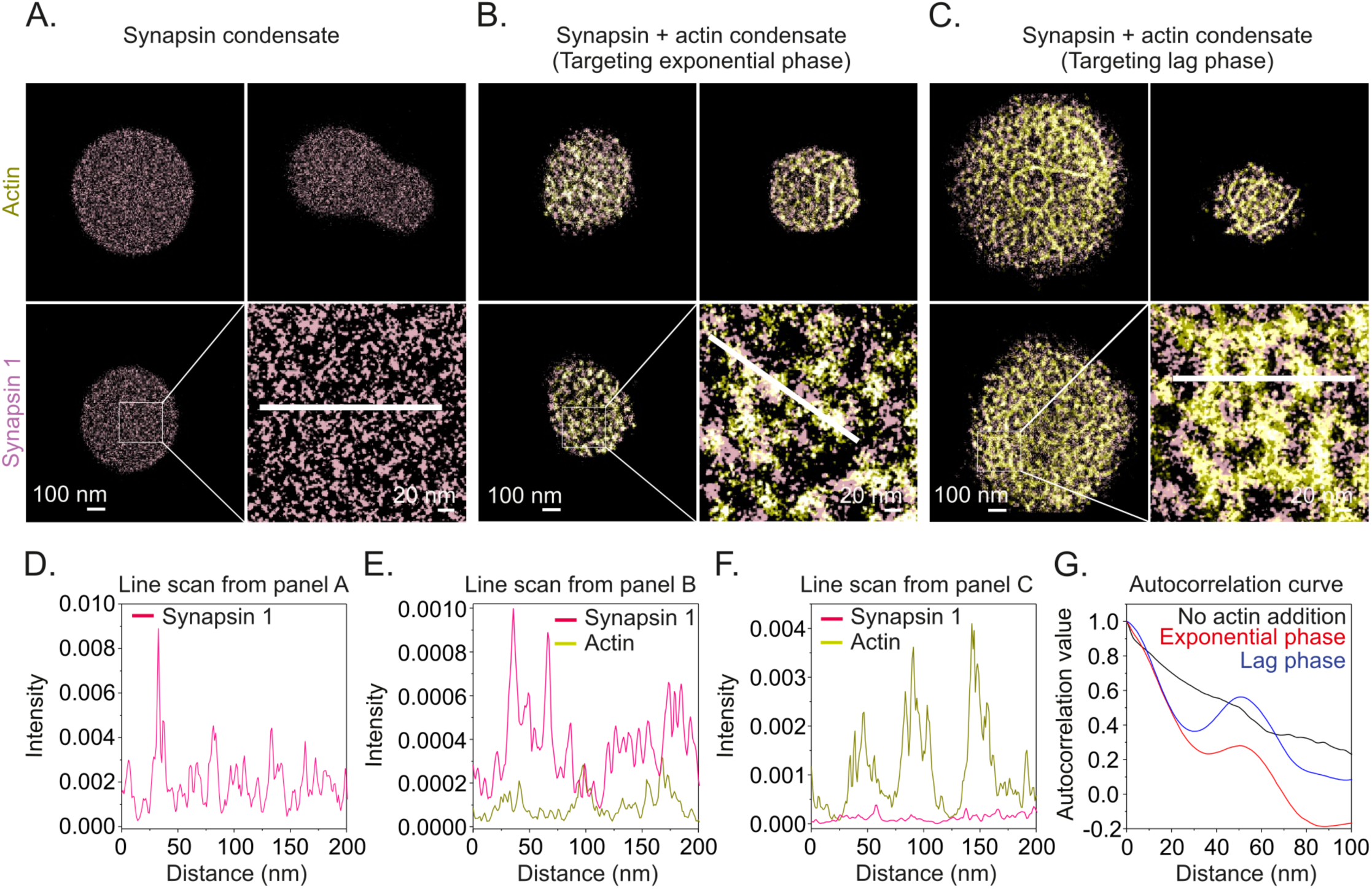
Regular synapsin-actin patterns observed *in vitro*. **A-C)** We generated synapsin-only condensates (panel **A**), or actin-synapsin condensates, composed of synapsin-1 and exponentially-growing actin filaments (panel **B**, to mimic the simultaneous addition of synapsin and actin filaments), or synapsin-1 and delayed-growth actin filaments (panel **C**, to mimic the addition of actin filaments to fully formed synapsin condensates). Images were taken by ONE microscopy. Scale bars = 100 nm, and 20 nm for the insets. **D-F)** To illustrate the composition of the condensates, we drew line scans across evident features (white lines in panels **A-C**). Repeated patterns are evident for actin. **G)** To detect repeated patterns, we analyzed the images using an autocorrelation approach. A repeated pattern with a periodicity of ∼50 nm is found for synapsin-actin condensates, while no such pattern is visible for synapsin-only condensates. *n* = 76, 64 and 33 condensates, respectively, for the conditions reported in panels **A-C**.

Both of these experiments led to the sequestration of actin within the condensates, as expected, consistent with the predictions of the simulations (**Fig. 2**). At the same time, visible actin filaments formed within the condensates (**Fig. 4B-C**; see image galleries in **Supplementary Fig. 10-11**). Further analysis of the structures revealed the existence of a periodic pattern, in the form of repeated dot-like actin structures in the synapsin-1 droplets (**Fig. 4E-F**). This pattern was also revealed by an autocorrelation analysis (**Fig. 4G**), which also confirmed that no clear organization can be detected for synapsin-only condensates. Overall, these experiments indicate that the combination of actin and synapsin-1 induce a repeated, periodic pattern *in vitro*.

### Simulations reveal that actin bundles are robust to synapsin droplet dissolution and reformation

Thus far, we have considered the interactions of a stable synapsin-1 droplet with actin. However, the synaptic vesicle cluster is a highly dynamic structure that is tightly regulated by upstream signaling events. PKA and CaMKII kinases phosphorylate the Ser9, Ser568, and Ser605 sites on synapsin-1, which causes a conformation change^16,45^ and reduces its affinity for actin^15,46,47^ and synaptic vesicles^27,48,49^. Phosphorylation of synapsin-1 thus leads to the dissolution of the droplet and release of synaptic vesicles within seconds^11,27^, mimicking the dynamics observed in hippocampal neurons in culture^49^. How does the dynamic dissolution of synapsin-1 condensates alter the actin structures observed above? To answer this question, we conducted simulations where synapsin droplets form, dissolve, and reform. To model synapsin-1 droplet dissolution, we tuned ε_Syn-Syn_ in our simulations. We first simulated synapsin and PEG as in **Figure 1B**, but we alternated ε_Syn-Syn_ between values where synapsin coalesced and where synapsin was diffuse to simulate initial droplet formation (ε_Syn-Syn_ = 2.0), droplet dissolution (ε_Syn-Syn_ = 1.0), and droplet reformation (ε_Syn-Syn_ = 2.0) over the time course of each simulation condition (**Fig. 5A**, **Supplementary Fig. 12A**). We tracked the normalized radius of gyration of synapsin over time to evaluate the progression of synapsin coalescence and dissolution (**Fig. 5B**, **Supplementary Fig. 12B**). We found that the average normalized radius of gyration initially decreased to ∼0.4–0.7 as synapsin particles coalesced (Frames 0–1000). However, once ε_Syn-Syn_ was reduced to 1.0, the synapsin particles rapidly diffused and the normalized radius of gyration returned to ∼1.0 (Frames 1000–2000). Upon increasing ε_Syn-Syn_ back to 2.0, the average normalized radius of gyration once again decreased to ∼0.4–0.7 as synapsin particles coalesced as before (Frames 2000–3000). Thus, we established a framework for reversible and dynamic modulation of synapsin droplet formation, reminiscent of experimental results obtained with synapsin-1 condensate dissolution^27^. Importantly, this simulation reveals that synapsin-1 droplets themselves do not contain any spatial or mechanical memory of their arrangement.

**Figure 5:**
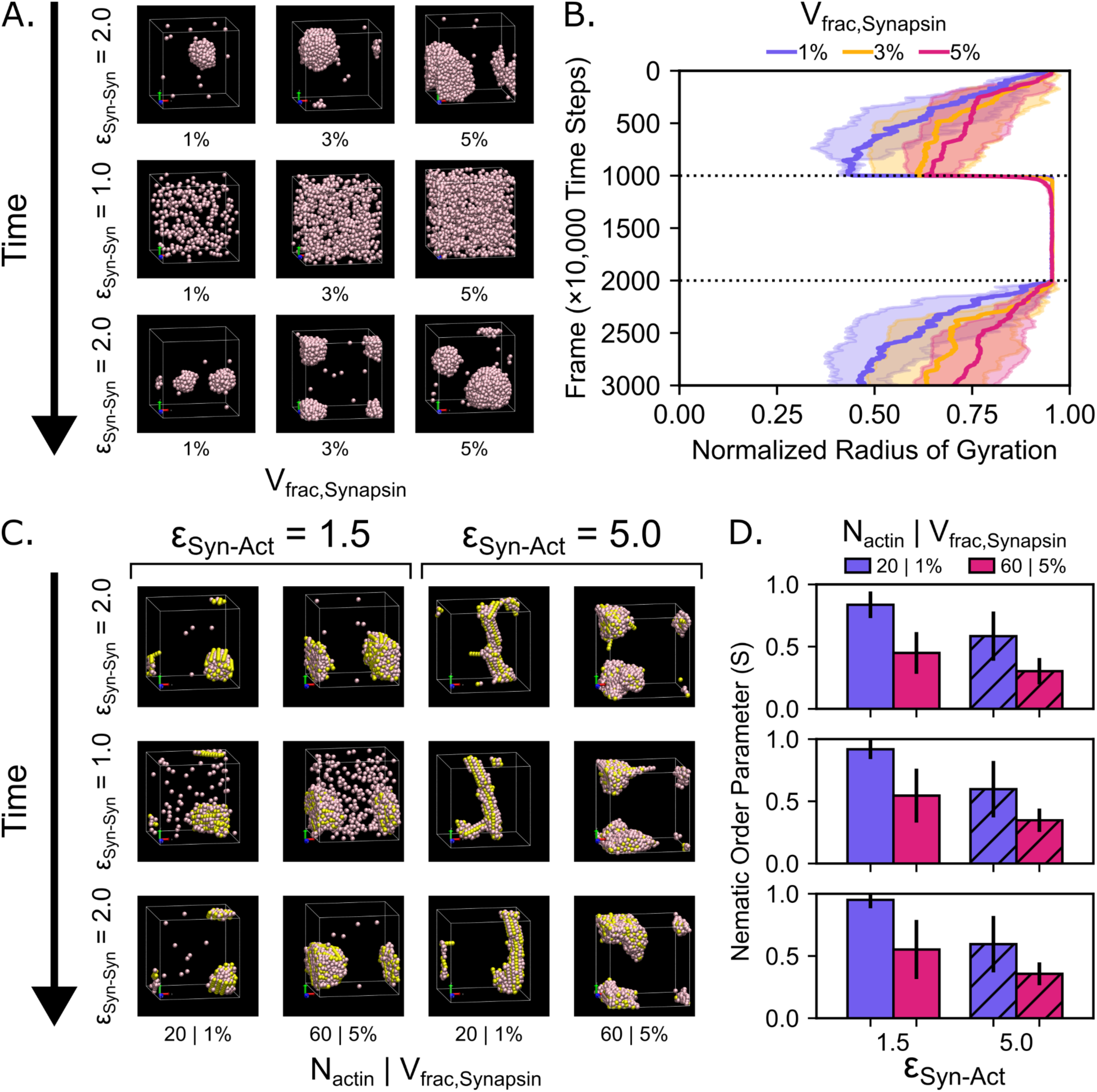
Altering the strength of Synapsin-Synapsin interactions can dissolve and reform synapsin droplets. **A)** Series of representative snapshots of simulations with synapsin and PEG particles depicting the initial formation, dissolution, and reformation of a synapsin droplet. ε_Syn-Syn_ is initially set to 2.0, lowered to 1.0 after frame 1000, and raised again to 2.0 after frame 2000. The volume fraction of synapsin is varied by column. The volume fraction of PEG is kept constant at 3%. PEG is not visualized in these snapshots. **B)** Time series showing the mean (solid line) and standard deviation (shaded area) of the normalized radius of gyration of the simulated synapsin particles, *n* = 10 replicates. The horizontal dotted lines indicate when ε_Syn-Syn_ was changed. **C)** Series of representative snapshots of simulations with the simultaneous addition of actin and synapsin. As with panel **A**, ε_Syn-Syn_ is initially set to 2.0, lowered to 1.0 after frame 1000, and raised again to 2.0 after frame 2000. The simulation condition is varied by column between (N_actin_ = 20, V_frac,Synapsin_ = 1%) and (N_actin_ = 60 and V_frac,Synapsin_ = 5%) for weak synapsin-actin attraction (ε_Syn-Act_ = 1.5) and strong synapsin-actin attraction (ε_Syn-Act_ = 5.0, used in all previous synapsin-actin simulations with fixed ε_Syn-Act_). The volume fraction of PEG is kept constant at 3%. PEG is not visualized in these snapshots. **D)** Bar chart depicting the mean nematic order parameter and comparing the simulation conditions in panel **C** at three illustrative time points (Frames 1000, 2000, and 3000) from the final frame of the three ε_Syn-Syn_ phases. Frame 1000 (top) corresponds to the end of the first phase, where ε_Syn-Syn_ = 2.0. Frame 2000 (middle) corresponds to the end of the second phase, where ε_Syn-Syn_ is decreased to 1.0. Frame 3000 (bottom) corresponds to the end of the third phase, where ε_Syn-Syn_ is increased back to 2.0. Empty bars correspond to ε_Syn-Act_ = 1.5, while diagonally striped bars correspond to ε_Syn-Act_ = 5.0. The error bars represent the standard deviation. *n* = 10 replicates, data from 10 time points in the last 100 frames of the simulations.

Next, we investigated how the dissolution and reformation of synapsin droplets would affect the arrangement of actin. We repeated the simulations of droplet formation-dissolution-reformation with actin present and tested the two exemplary conditions from previous simulations, (N_actin_ = 20, V_frac,Synapsin_ = 1%) and (N_actin_ = 60 and V_frac,Synapsin_ = 5%), in the cases of weak synapsin-actin attraction (ε_Syn-Act_ = 1.5) and strong synapsin-actin attraction (ε_Syn-Act_ = 5.0), for a total of four different simulation conditions (**Fig. 5C**). Collectively, these conditions capture the range of nematic order seen in all our simulations. For additional conditions, please see **Supplementary Figure 13**. We found that actin structures retained their organizational features even when the synapsin droplet was dissolved (**Fig. 5C-D**). These results suggest the possibility that once an actin scaffold is templated by the synapsin droplet, it is able to maintain its organization even when the droplet is dissolved and reformed, hinting at a possible spatial or mechanical memory.

### Experiments reveal that dissolving synapsin condensates leaves actin largely unchanged

To test the effects of condensate dissolution *in vitro*, we sought to preferentially disrupt the synapsin phase, without directly influencing actin. We relied on synapsin-actin condensates, incubated for 30 minutes, to enable the formation of actin filaments, which were treated with the aliphatic alcohol 1,6-hexanediol (10% w/v), which is known to disrupt hydrophobic interactions^50^, and perturbs LLPS-based arrangements, including synapsin-actin condensates^11^. Treatment with 1,6-hexanediol strongly reduced the synapsin signal intensity (**Fig. 6A**), but only a minor proportion of the actin signal was lost (**Fig. 6B-C**). All of the morphological parameters of the actin arrangements remained unchanged after this treatment (**Fig. 6D-I**), confirming the suggestion that, once formed, the actin arrangements induced by synapsin condensates are stable, even without the support of synapsin.

**Figure 6:**
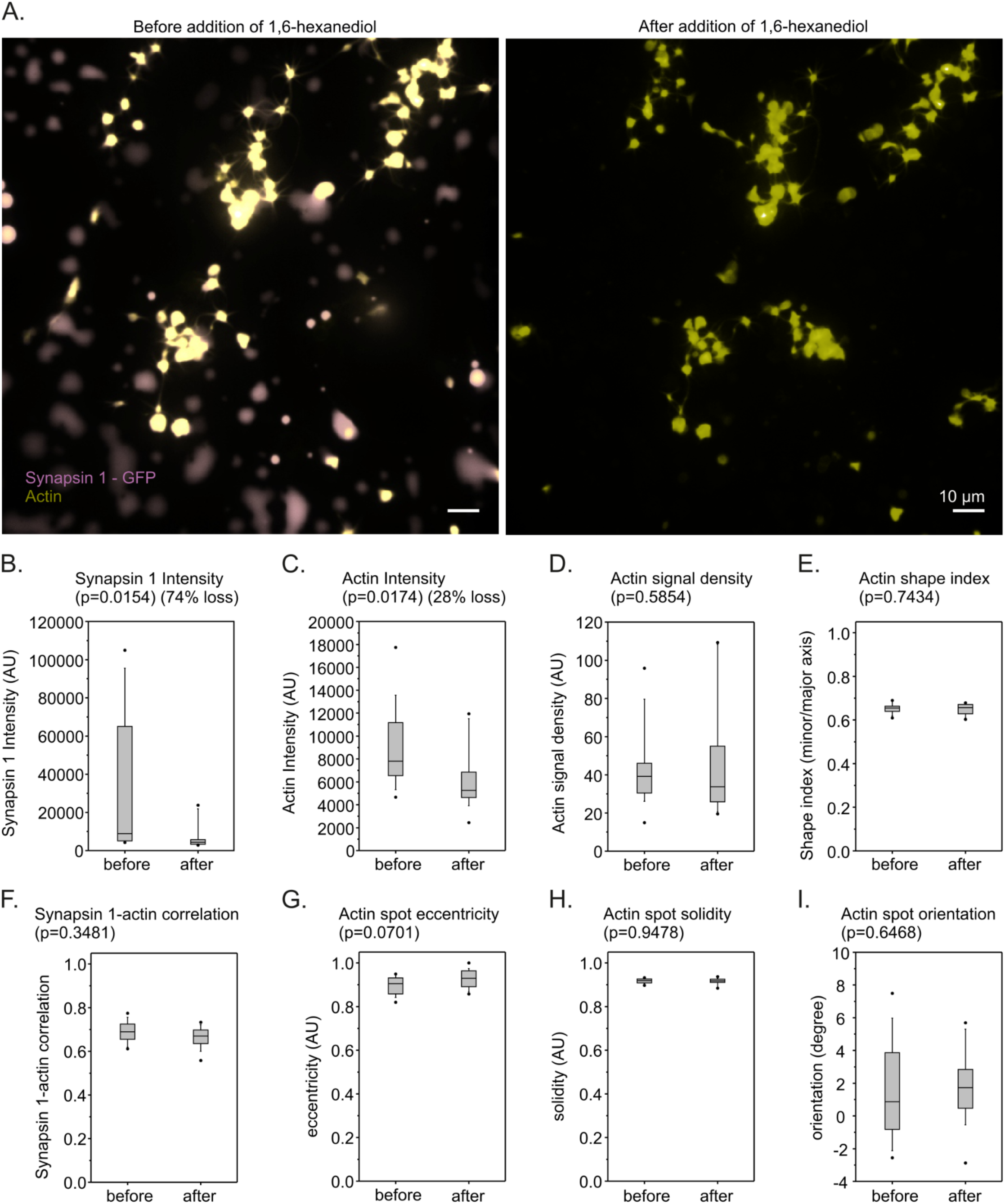
Disruption of synapsin condensates *in vitro* leaves actin organization largely unaffected. **A)** Synapsin-actin condensates were imaged before (left) and after (right) the addition of 1,6-hexanediol (10% w/v). Scale bars = 10 µm. **B-C)** The intensity of synapsin in the condensates was strongly reduced, with only around 26% of the synapsin persisting, while the actin intensity was only lowered by ∼28%. **D-I)** We analyzed several parameters of the actin arrangements, including morphological features (as the shape index, the spot orientation, spot solidity and spot eccentricity), intensity features (the signal density) or the correlation between actin and the remaining synapsin. No significant differences could be found between the conditions before or after 1,6-hexanediol treatments (Mann-Whitney rank sum tests, p values indicated in the respective panels). *n* = 14-15 imaging areas for each condition.

## Discussion

In this study, we used a combination of physics-based simulations and *in vitro* reconstitution experiments to shed light on the nanoscale organization of actin filaments in synapsin droplets. Our simulations revealed a number of fundamental aspects. First, the actin-synapsin composite environment drives the emergence of nematic structures (**Fig. 2**). More importantly, the nematic organization of actin filaments in synapsin droplets leads to the emergence of a regularly organized structure that is a function of actin and synapsin interaction parameters (**Fig. 3**). Independently, nanoscale imaging revealed a repeated actin pattern with a periodicity of ∼50 nm in reconstituted synapsin condensates (**Fig. 4**). Simulations showed that actin organization persisted through cycles of droplet dissolution and reformation (**Fig. 5**), while experiments demonstrated that actin architecture is preserved following condensate disruption (**Fig. 6**). Together, our results suggest that actin structures in synapsin droplets may template a form of structural or mechanical memory in synapses.

To our knowledge, this work is distinct from previous work on actin organization in phase separated droplets, where we and others have contributed extensively. At the mesoscale, experimental and computational studies of actin organization in droplets formed from actin-binding proteins have revealed bundled structures such as rings, discs, rods, asters, and shells^6–10,51^. Complementary studies have shown how the surface-induced phase separation at lipid bilayers promotes localized bundle formation^52,53^, how spatial confinement affects the viscoelastic properties of semiflexible polymer networks of actin filaments^54^, and how the molecular properties of actin crosslinkers determine the viscoelastic properties and drive the dynamic remodeling of actin networks^55^. Further work has shown that membrane tension drives cortical actin reorganization^56^ and that crosslinked actin bundles exhibit liquid-like properties^57^. While previous studies focused on how actin crosslinking, membrane interactions, and spatial confinement drive mesoscale network architecture and dynamics, the present study examines the nanoscale organization of actin within synapsin condensates. By combining coarse-grained simulations with nanoscale microscopy, we demonstrate that synapsin-actin interactions drive the emergence of a periodic nanoscale actin architecture that remains resilient through cycles of droplet dissolution and reformation. The persistence of these actin scaffolds suggests a physical mechanism for synapse-specific structural memory that enables local synaptic architecture to survive continuous activity, condensate turnover, and plastic remodeling. This view aligns with observations of actin-based memory in systems that range from the postsynaptic compartment to migrating cells^58,59^.

While our findings provide key insights into synapsin-actin organization within the synaptic vesicle cluster, we acknowledge the limitations of using coarse-grained particles with isotropic LJ potentials for our model. While this approach abstracts more complex biological features, this model is capable of capturing the minimal physics that govern the system. To build upon these simulations, future models could incorporate dynamic filament nucleation and polymerization, increased filament length and flexibility, electrostatic particle interactions, and explicit binding sites that better capture finer molecular interactions.

Our findings have particular implications for synaptic function. Actin has been studied extensively in the context of presynaptic function, especially in the presynaptic bouton where it forms filaments that are visible within the vesicle clusters^28,60,61^, albeit most actin is localized in a meshwork that surrounds the clusters that is especially evident in large synapses as the neuromuscular junctions^62–64^. The actin filaments surrounding the clusters may be used in stabilizing the liquid phase of the cluster^65^, but they have also been known for decades to participate in endocytosis, retrieving synaptic vesicles from the peri-active zone, and bringing them to the vesicle cluster^66^. Actin at the periphery of the vesicle cluster also assists with the membrane invagination process during endocytosis, through the action of myosin motors that depend on the presence of peri-active zone actin filaments^67^.

Since actin dynamics are an essential component of synaptic physiology, and especially of endocytosis, it has long been hypothesized that actin forms a scaffold that enables the organization of the synaptic vesicles^68^. Alternatively, the vesicles may organize without an explicit need for actin, by phase separation^69^, with actin recruitment to the vesicle clusters being a secondary phenomenon. Actin seems to accumulate not only on vesicle clusters, but also around patches of vesicle proteins on the plasma membrane^70,71^, suggesting that vesicle proteins are able to recruit actin, irrespective of where they find themselves, which would argue for the second model (phase separation). Crucially, modulation of synapsin, by use of antibodies^72,73^ disorganizes the vesicle clusters, suggesting that they are maintained by vesicle-synapsin interactions and phase separation rather than by actin scaffolding. This implies that, overall, the actin scaffold model, in which actin organizes the vesicle cluster, is currently difficult to accept as the dominant mechanism^73^, and that the alternative hypothesis, that the vesicle cluster organizes the actin cytoskeleton, primarily through interactions between vesicle-associated synapsin molecules and actin, better fits current evidence.

The function of the actin cytoskeleton within the vesicle cluster remains challenging to interpret due, in no small part, to the fact that these filaments have been difficult to observe in imaging experiments. They are unlikely to serve as routes for vesicle transport towards the active zone, since most synaptic vesicle movement within boutons is not directed, but diffusive, thus being incompatible with actin-based transport^74–76^. Moreover, actin organization at the presynapse has to accommodate concomitant processes where distinct actin morphologies, such as actin tracks, interface shells, and inter-vesicular connectors, need to sustain repeated rounds of SV release^61,77^. These dynamic structural networks adapt continuously to support vesicle mobilization and active zone cytomatrix rearrangements during high-frequency neurotransmission. Our experiments suggest that this cluster-internal meshwork is involved in forming and stabilizing the core of the synaptic vesicle cluster. In principle, vesicles need to remain mobile within the cluster core to maintain their ability to participate in functional reactions upon demand. At the same time, the vesicle cluster needs to remain stable to maintain a large pool of soluble cofactor proteins, without which exo- or endocytosis reactions would be impossible^78^. To ensure that these two contrasting needs are fulfilled, the actin cytoskeleton stabilizes the vesicle cluster core. Actin would ensure against functional changes triggered by periods of high synaptic activity, which result in synapsin phosphorylation and vesicle diffusion. Even if vesicles are lost from boutons during such activity conditions, the actin core could possibly remain in place, ready to collect again the vesicles during the subsequent resting periods. This actin-induced stability of the vesicle cluster would enable it to provide a certain memory to the synapse, and to control its functional parameters over time^79^.

Our imaging experiments provide a first view of an organized, periodic pattern of actin filaments within a condensate. In principle, the components of condensates should be mobile and lack any inherent special structure, as was the case for the synapsin-only condensates. However, adding actin filaments to the system resulted in a regularly organized pattern. Our investigation leads to the hypothesis that condensates containing filamentous objects, including cytoskeletal components and even RNA molecules, may have distinct nanoscale structures that remain to be discovered. Importantly, such structures may modulate both the function and dysfunction of proteins within condensates. These structures potentially include aggregated forms of alpha-synuclein, a prominent protein in Parkinson’s Disease and a known component of several condensates including P-bodies^80^ and synapsin condensates^25,81^. Furthermore, recent studies have revealed that synapsin-1 condensates contain interfacial electric potentials^82^ and catalyze specific chemical transformations unattainable in dilute phases^83^. Thus, the specialized internal microenvironment of synapsin-1 condensates may allow for the distinct diffusive behaviors of synaptic vesicles within clusters and can also support actin polymerization. Along these lines, our computational discovery and experimental validation of actin nanostructures within synapsin-1 suggests that the combination of actin filaments and biomolecular condensates may generate functional reaction centers essential for high fidelity neural transmission.

## Methods

### Model Development

We constructed a mesoscopic model that represents synapsin, actin, and PEG with spherical LJ particles in LAMMPS (**Fig. 1A**), a widely used molecular dynamics program^39^. We use reduced Lennard-Jones units for our LAMMPS model, so length scales, energy, mass, and simulation time are all expressed in a dimensionless form relative to fundamental reference units for length (σ), energy (ε), mass (*m*), and time (**τ**). Synapsin is modeled as a single spherical particle that functionally represents a monomer that is attracted to actin filaments and to other synapsin particles. Performed actin filaments are modeled as semi-flexible linear polymers that consist of a string of 10 spherical particles subjected to bonded interactions with harmonic stretching and bending potentials that match experimental values, as previously described^8^. The PEG is modeled as a single spherical particle that functions as a crowding agent with solely repulsive steric interactions with other species.

To characterize the effective mass and diameter of each particle in the system without introducing atomic complexity to our model, the reduced mass (*m*) and steric particle diameter (σ) parameters were assigned based on molecular weights (M) and effective spatial footprints of each species. We chose full-length synapsin-1 (M ≈ 74 kDa, UniProt P17600^84^) to serve as the reference species (m_synapsin_ = 1.0*m*, σ_Syn-Syn_ = 1.0σ). Synapsin-1 consists of both ordered and disordered domains, which means that the steric volume of synapsin can vary depending on the solution. Previous experimental measurements for the radius of gyration of synapsin indicated that the protein was partially unfolded and yielded a value of 4.2 nm^85^, although synapsin may adopt a different radius in the dense phase of a condensate. Individual actin monomers (M ≈ 42 kDa, UniProt P60709^84^), m_actin_ = 42/72 ≈ 0.567*m*) within actin filaments are about 5.5 nm in length while actin filaments have a cross-sectional diameter of about 7-9 nm^86,87^. Since synapsin and actin occupy similar steric envelopes, we decided to coarse-grain both particles as single beads with the same steric diameter (σ_Syn-Syn_ = σ_Act-Act_ = 1.0σ). We chose to base the PEG crowder on PEG8000 (M ≈ 8 kDa, m_PEG_ = 8/72 ≈ 0.108*m*) and used calculated radii of gyration from experimental measurements^88^ to determine its steric particle diameter (σ_PEG-PEG_ ≈ 0.8σ). The characteristic length for the interactions between particle types (σ_ij_) were subsequently determined by the arithmetic mean of the two steric particle diameters.

All non-bonded interactions between particles are governed by the Lennard-Jones potential:

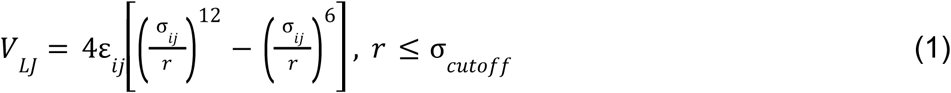

Attractive particle interactions, specifically synapsin-synapsin and synapsin-actin interactions, use a standard cutoff distance of σ_cutoff_ = 2.5σ. Meanwhile, particle interactions that are purely repulsive, specifically actin-actin interactions and any interaction pair that involves PEG crowders, use a cutoff distance of σ_cutoff_ = 2^1/6^σ. Since the potential V_LJ_ reaches a minimum ε value at r = 2^1/6^σ, by setting σ_cutoff_ = 2^1/6^σ, we can ensure that only the repulsive regime remains.

### LAMMPS Simulation Protocol

The simulation space consisted of a cubic box with side lengths of 25σ under periodic boundary conditions in all directions (x, y, and z). Particles are randomly placed into the simulation space using distinct random number seeds. Energy minimization was conducted using the default conjugate gradient algorithm to resolve any initial steric overlaps. Minimization proceeded until either the energy tolerance reached 10^-4^ or the force tolerance reached 10^-6^ within 10,000 iterative steps or 100,000 force or energy evaluations. After energy minimization, initial velocities for the particles were generated from a Gaussian distribution corresponding to the reduced target temperature T* = k_B_T/ε = 1.0 using a random number seed. Simulations were performed using an NVT-like canonical ensemble at T* = 1.0 using Langevin dynamics, which is achieved by combining NVE time integration with a Langevin thermostat configured with a dampening parameter of 1.0 and a random number seed. The equations of motion were integrated using the discrete Verlet algorithm with a discrete time step of 0.0005. Neighbor lists were updated every 10 time steps with a skin distance of 0.3σ. System trajectories were sampled every 10,000 time steps. Production runs were performed over 30,000,000 time steps in all conditions except for simulations where actin is added to the system after synapsin, in which case the production runs were performed over 40,000,000 time steps. Please refer to **Supplementary Table 1** for a detailed list of parameters used. The LAMMPS simulations were performed on the Triton Shared Computing Cluster (TSCC) at the San Diego Supercomputer Center (SDSC)^40^.

## Experimental Methods

### Actin Stock Preparation

Unlabeled actin (Cytoskeleton, #AKL99A) and biotinylated actin (Cytoskeleton, #AB07A) stock solutions were prepared by reconstitution in ultrapure water according to the supplier’s instructions and dilution in 1x PBS supplemented with 0.2 mM ATP to a concentration of 2 mg/mL.

### Expression and Purification of EGFP-Synapsin 1

6xHis-EGFP-rnSynapsin 1 was expressed and purified at 4 °C following previously described protocols^25–27^. Briefly, 6xHis-EGFP-rnSynapsin-1 was expressed in Expi293F™ cells (ThermoFisher) for 3 days after induction. Cells were further lysed with three freeze-thaw cycles, consisting of rapid freezing in liquid nitrogen followed by thawing at 37 °C in buffer A (25 mM Tris-HCl (pH 7.4), 300 mM NaCl, 25 mM imidazole and 0.5 mM TCEP) supplemented with EDTA-free Roche Complete protease inhibitors, 10 µg/mL DNase I, and 1 mM MgCl_2_. The lysate was then clarified by centrifugation for 1 h at 20,000 × g. The resulting soluble supernatant was then added on a Ni-NTA affinity column (HisTrap™HP, Cytiva), and purification was carried out on a ÄKTA pure 25 M system. The Ni-NTA column was further washed with buffer A containing 40 mM imidazole and finally the bound protein was eluted using 400 mM imidazole. Fractions obtained from affinity purification were concentrated using a 30 K MWCO protein concentrator (Pierce) and were further purified by size-exclusion chromatography with Superdex™ 200 Increase 10/300 column (GE Healthcare) equilibrated in 25 mM Tris-HCl (pH 7.4), 150 mM NaCl, and 0.5 mM TCEP. Elution fractions were evaluated by SDS-PAGE. Purified synapsin-1 was then snap-frozen in liquid nitrogen and stored at −80 °C until further use.

### Preparation of Actin-Condensate Samples

To analyze the structural interplay between actin filaments and EGFP-synapsin 1, samples were prepared using three assembly pathways, as described below.

### Reagent and Solution Formulations

#### Labelled EGFP-synapsin 1 complex

To prepare the labeled protein complex, 0.75 µL of 40 µM EGFP-synapsin 1 was incubated with 6 µL of 5 µM Cy3-conjugated anti-GFP nanobody (Nanotag, #N0303-SC3-L) in 12 µL of 1× PBS supplemented with 2 mM TCEP. The mixture was incubated for 30 min at room temperature to allow nanobody binding.

#### Condensate mixture

Macromolecular crowding and subsequent condensation were induced by adding 12 µL of 25% polyethylene glycol 8000 (Sigma-Aldrich, 86101) to the labelled EGFP-synapsin 1 complex, followed by gentle mixing. The mixture was deposited onto a coverslip and incubated for 60 min at room temperature.

#### EGFP-synapsin 1 monomer control

To maintain the same overall concentration of synapsin 1 as in the condensate condition, 12 µL of 1× PBS was added to the labelled EGFP-synapsin 1 complex, followed by gentle mixing. The mixture was then deposited onto a coverslip.

#### Unlabelled actin working solution

Unlabelled actin working solution at a protein concentration of 0.2 mg/mL was prepared by mixing 5 µL of unlabelled actin stock solution with 45 µL of 1x PBS supplemented with 0.2 mM ATP and 0.5 mM DTT.

#### Biotinylated actin working solution

Biotinylated actin working solution at a protein concentration of 0.2 mg/mL was prepared by mixing 2 µL of biotinylated actin stock solution with 18 µL of 1x PBS supplemented with 0.2 mM ATP and 0.5 mM DTT.

#### Actin monomer pool

The unlabelled and biotinylated actin working solutions were mixed at a 1:4 volumetric ratio, yielding an actin monomer pool containing 80% biotinylated actin. Fully biotinylated actin did not polymerize efficiently under these conditions. To modulate the initial polymerization kinetics before assembly, the actin monomer pool was incubated for 30 min either at room temperature, to favor an accelerated exponential phase, or on ice, to favor a prolonged lag phase. At this ATP concentration, macroscopic filament assembly remained suppressed before addition of the polymerization cocktail.

#### Polymerization cocktail

Actin filament assembly was initiated by mixing 75 µL of 1x PBS with 0.2 mM ATP, 10 µL of 1x PBS with 10 mM ATP, and 15 µL of actin monomer pool.

### Assembly Pathways

#### Condensate plus actin polymer

A 20 µL aliquot of the polymerization cocktail was added directly to coverslips containing pre-incubated condensates, followed by incubation for 3 h at room temperature. This condition included two subconditions in which actin polymerization was biased either toward an accelerated exponential phase or toward a prolonged lag phase, as described above.

#### Synapsin 1 monomer plus actin polymer

A 20 µL aliquot of the polymerization cocktail was added directly to coverslips containing EGFP-synapsin 1 monomer control samples, followed by incubation for 3 h at room temperature.

#### Condensates without actin

Coverslips containing EGFP-synapsin 1 condensates pre-incubated for 60 min at room temperature were processed without addition of actin.

### Chemical Stabilization and Fluorescent Labelling

To preserve structural organization during downstream processing, actin-synapsin 1 assemblies were chemically stabilized on coverslips before expansion.

#### Filament stabilization

Actin filaments were stabilized by adding 15 µL of 0.3 mM phalloidin (Thermo Fisher Scientific, P3457) prepared in F-buffer, followed by incubation for 15 min at room temperature.

#### Functional anchoring

Samples were anchored by incubation with a freshly prepared mixture containing 7.5 µL of 1× PBS and 2.5 µL of 10 mg/mL acryloyl-X SE (AcX; Thermo Fisher Scientific, A20770) for 30 min at room temperature.

#### Chemical fixation

Samples were fixed by adding 15 µL of 1.5% methacrolein (Sigma-Aldrich, 133035) in 1× PBS to obtain a final concentration of 0.25%, followed by incubation for 15 min at room temperature.

#### Biotin labelling

Biotinylated actin polymers were labelled by adding 20 µL of diluted streptavidin-STAR635P (Sigma-Aldrich, 04683), prepared by adding 1.1 µL of a 1 mg/mL stock solution to 100 µL of 1× PBS. After incubation for 45 min at room temperature, coverslips were washed three times with 1× PBS to remove unbound fluorophores before gelation. For condensate-only samples, the phalloidin stabilization and streptavidin labelling steps were omitted, and 12.5 µL of methacrolein solution was used for fixation.

### Kinetic Modulation of Actin-Synapsin 1 Assembly

The EGFP-synapsin 1 and nanobody complex was prepared by incubating 0.75 µL of 40 µM EGFP-synapsin 1 with 6 µL of 5 µM Cy3-conjugated anti-GFP nanobody in 12 µL of 1× PBS for 30 min at room temperature. Condensation was then induced by adding 12 µL of 25% PEG in 1× PBS. The resulting 30 µL mixture was transferred onto a coverslip and incubated for 60 min to induce phase separation.

To systematically control the initial polymerization kinetics during actin-synapsin 1 assembly, the actin monomer pool, prepared by mixing unlabelled and biotinylated actin working solutions at a 1:4 volumetric ratio (∼82% biotinylated actin), was distributed into two distinct 30-min thermal pre-incubation regimes. To favor an accelerated exponential polymerization phase, the monomer pool was incubated at room temperature, whereas to induce a prolonged lag phase, the pool was incubated on ice. Following this 30-min kinetic modulation step, 15 µL of the respective monomer pool was mixed with 75 µL of G-buffer and 10 µL of F-buffer (10 mM ATP) to form the active polymerization cocktail. Finally, to start the assembly, we added 20 µL of our prepared actin polymerization cocktail directly to the 30 µL of pre-incubated synapsin 1 condensates. This combined 50 µL mixture was incubated for 3 hours at room temperature before we proceeded with chemical stabilization.

### X10 Expansion

Samples were anchored to the hydrogel using a combination of acryloyl-X SE and methacrolein, as described above. Methacrolein served both as a fixative and as an anchoring reagent^89^. The gel monomer solution was prepared as described previously. An 80 µL drop of monomer solution was deposited onto Parafilm and carefully brought into contact with the inverted coverslip carrying the sample. Gels were polymerized overnight at room temperature in a dark, humidified chamber.

Sample homogenization was performed as previously described^37,90,91^. In brief, gels were incubated overnight at 37 °C with 8 U/mL proteinase K (Merck, P4850) in digestion buffer containing 800 mM guanidine HCl, 2 mM CaCl_2_ and 0.5% Triton X-100 in 50 mM Tris buffer. After digestion, isotropic physical expansion was achieved by successive washes in excess double-distilled water over several hours until the gels reached full expansion. Fully expanded gels reached final diameters of 15-17 cm and were stored in water-filled chambers until imaging. Before data acquisition, gels were cut into rectangular pieces of suitable size and transferred to the imaging chamber. A detailed step-by-step sample preparation protocol, including a visual workflow, has been described previously^37,91^.

### Fluorescence Imaging

Initial assessment of non-expanded samples was performed on an Olympus IX71 epifluorescence microscope equipped with a 100×, 1.41 NA oil-immersion objective. Images were acquired with an F-view II CCD camera, corresponding to a pixel size of 64.5 nm.

ONE microscopy imaging of expanded samples was performed as previously described^37,92–95^. Briefly, samples were imaged using a Leica TCS SP5 STED microscope operated in confocal mode. Samples were excited with a 561 nm laser for Cy3 or a 633 nm laser for STAR635P through an HCX Plan Apochromat 100×, 1.4 NA oil-immersion STED objective. Fluorescence emission was collected with a PMT detector for Cy3 and a HyD detector for STAR635P. Wavelength selection was controlled by an acousto-optic tunable filter. Images were acquired at a line-scanning speed of 8 kHz. Time-series movies consisting of 1,500 frames were acquired in 8-bit format with an image size of 128 × 128 pixels, corresponding to a calibrated pixel size of 98 nm. Final high-resolution images were reconstructed from raw movies in Fiji using the ONE plugin^37,96^.

### Image Autocorrelation Analysis

Autocorrelation analysis of condensates was performed using custom MATLAB scripts (R2023b; MathWorks). Rectangular regions of 225 × 225 nm were manually selected in each analysed image and centred on synapsin 1 or actin signals. Images were then rotated to align the signals to a rectangular spot positioned 50 nm from the image centre, ensuring that all patterns had the same approximate orientation. The aligned images were averaged, and autocorrelation curves were calculated from the averaged images.

For the analysis of live-imaging experiments, synapsin 1- and actin-containing regions were identified using automated thresholding, and their features were extracted using the MATLAB regionprops function. Statistics were calculated from averages of individual imaging experiments rather than from values of individual condensates.

### Chemical Disruption of Synapsin 1 Condensates

Synapsin 1 condensates were generated by incubating 10 µM EGFP-synapsin 1 with 7.5% (w/v) PEG 8000 in a reaction buffer containing 25 mM Tris-HCl (pH 7.4), 150 mM NaCl, and 0.5 mM TCEP for 15 min. The resulting preformed condensates were then added from above to electroformed DOPC GUVs that had been pre-loaded onto a 15-well glass slide (ibidi, 81506). Condensates were allowed to settle on the GUV surface for approximately 20 min. ATP- and magnesium-exchanged actin was subsequently added from the top to the condensate-GUV mixture. For visualization, unlabeled actin (Hypermol, 8101-01) was supplemented with ATTO647-labeled actin (8158-02). The resulting final reaction mixture contained 3% (w/v) PEG 8000, 4 µM Syn1-FL, 4 µM actin, 74.5 mM NaCl, 0.5 mM ATP, and 150 mM sorbitol. Following actin polymerization for approximately 30 min, 1,6-hexanediol was added to a final concentration of 10%. The sample was gently mixed and incubated for an additional 5 min. Images were acquired at the same locations before and after the addition of 1,6-hexanediol. Imaging was performed on a Nikon spinning-disk confocal CSU-X (SDC CSU-X) microscope equipped with two EMCCD cameras (iXon3 DU-888 Ultra), Andor Revolution SD System (CSU-X), and a PL APO163 60x/1.4 NA, oil immersion objective. Excitation wavelengths were: 488 nm for EGFP-synapsin 1 and 647 nm for ATTO647-labeled actin. The data were originally acquired for a published project^11^. In the present study, the original data were reanalyzed to assess the stability of actin arrangements following the disruption of synapsin 1 condensates.

## Data Analysis

### Radius of Gyration

The radius of gyration R_g_ is directly calculated from the pairwise minimum-image distances between synapsin particles in order to account for the periodic boundary conditions. A distance matrix is populated and then the squared radius of gyration is evaluated over all distinct pairwise combinations of synapsin particles using the following formula:

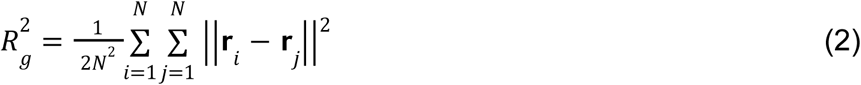

where **r**_i_ and **r**_j_ are the 3D coordinates of particles i and j, N is the number of synapsin particles, and we evaluate 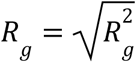 to calculate the radius of gyration. We then identify the global maximum measured R_g_ value across all conditions and their replicates and divide each measured R_g_ by the maximum R_g_ to calculate the normalized radius of gyration, ensuring values bounded by 0 and 1.

### Nematic Order Parameter

To quantify actin filament orientational alignment, we construct the symmetric nematic order tensor **Q**:

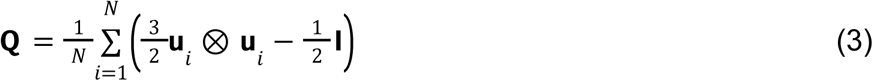

where **u**_i_ is the unit orientation vector for each actin filament, N is the total number of actin filaments, and **I** is the identity matrix. We then diagonalize **Q** and determine the eigenvalues and eigenvectors, **Qn** = *S***n**. The scalar nematic order parameter S corresponds to the maximum eigenvalue of **Q**:

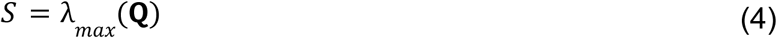

while the nematic director **n** is the associated eigenvector.

### Statistics and Reproducibility

Due to the stochastic nature of our simulations, we performed 10 replicate simulations per condition to yield a sufficiently large sample size for statistical analyses of the results. All LAMMPS trajectories initialized particles with a random distribution of initial positions and initial velocities. All data analysis for simulations was performed on Python 3.11.9.

The simulation data presented in **Figure 2** and **Supplementary Figure 2** required careful treatment to extract the statistical analysis shown in **Supplementary Figure 3** and **Supplementary Table 2**. Since the scalar nematic order parameter is bounded to the domain *S* ∈ [0, 1] in our system, we transform the data using the logit function:

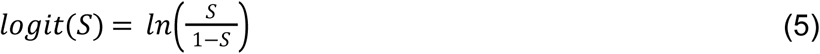

which maps S values onto an unbounded domain to homogenize the resulting variance. Model assumptions were then tested for the transformed residuals. Even with the logit transformation, Levene’s test for homoscedasticity revealed a difference in variances between groups (W = 1.5814, *P* = 0.0095). Normality was evaluated using the Shapiro-Wilk test (W = 0.9818, *P* < 0.0001) and shape parameters skewness and kurtosis. Although this test indicated a statistically significant difference from ideal normality, this difference may be due to the high sensitivity of the Shapiro-Wilk test^97^ for our relatively large sample size (*n* = 500). Furthermore, the logit-transformed data exhibited minimal skewness (0.2418) and minimal excess kurtosis (1.5691), which are within tolerance limits for linear model validity.

A three-way Analysis of Variance (ANOVA) was performed on the full logit-transformed nematic order dataset (*n* = 500) to evaluate the main effects and interactions of the number of actin filaments (N_actin_), the volume fraction of synapsin (V_frac,Synapsin_), and the order of actin addition (Order) (**Supplementary Table 2**). The three-way ANOVA revealed significant main effects for N_actin_ (F = 68.8051, *P* < 0.0001), V_frac,Synapsin_ (F = 9.3181, *P* < 0.0001), and Order (F = 104.6974, *P* < 0.0001), and significant two-way interactions between N_actin_ and Order (F = 5.7717, *P* = 0.0002) and V_frac,Synapsin_ and Order (F = 2.7409, *P* = 0.0282). Additionally, the three-way ANOVA also revealed that the two-way interaction between N_actin_ and V_frac,Synapsin_ (F = 1.2368, *P* = 0.2357) and a the three-way interaction between N_actin_, V_frac,Synapsin_, and Order (F = 1.2427, *P* = 0.2314) are not statistically significant. These results confirm that the number of actin filaments and the order of actin addition, both their main effects individually and the two-way interaction between N_actin_ and Order, produce statistically significant differences in nematic order.

Based on the ANOVA results, the logit-transformed dataset was partitioned by the order of actin addition (*n* = 250 for each Order condition), and individual simulation replicates for each N_actin_ condition were then pooled across all V_frac,Synapsin_ conditions (*n* = 50 for each N_actin_ condition). Pairwise comparisons between N_actin_ conditions within each order of actin addition condition were then evaluated using Games-Howell post-hoc tests on these datasets. Games-Howell was selected because it adjusts the degrees of freedom to account for unequal variances without assuming homoscedasticity. Additionally, Games-Howell is robust to mild non-normality for relatively large sample sizes per group^98^, such as in our analysis (*n* = 50). These comparisons allowed us to evaluate the effect of N_actin_ when actin was added simultaneously with synapsin (**Supplementary Fig. 3A**) and when actin was added after synapsin (**Supplementary Fig. 3B**). The means and 95% confidence intervals calculated for the logit-transformed data in **Supplementary Figure 3** were reverse-transformed using the expit function (logistic function):

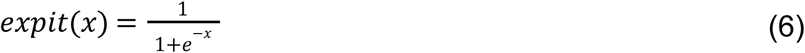

All statistical analysis on simulation data was performed using the scipy.stats, pingouin, and statsmodels Python packages.

For biological experiments, tens to hundreds of condensates were analyzed for each experiment (*n* values indicated in the respective figure legends). In **Figure 6**, Mann-Whitney rank sum tests were used to test significant differences between conditions; the *n* value here refers to imaging areas, which typically contain more than 10 condensates per area. No statistical tests were employed in **Figure 4** and **Supplementary Figure 8**.

## Data Availability

The LAMMPS input files and data generated in this study from experiments and simulations will be made available on publication.

## Code Availability

Python scripts used to analyze LAMMPS trajectories will be made available on publication.

## Acknowledgements

This research was supported by the National Science Foundation (NSF) through MODULUS Grant MCB 2327243 to P.R., the National Institutes of Health (NIH) through the Molecular Biophysics Training Grant T32 GM139795 to D. Mansour, the European Research Council (ERC) Grant (101078172 to D. Milovanovic), and by German Research Foundation (DFG) grants to S.O.R. and S.K. (SFB1286/B02) and D. Milovanovic (SFB1286/B10). We thank Dr. Emmet Francis for their guidance on statistical analysis and Prof. Christopher T. Lee for discussions and their advice on LAMMPS simulations and metrics for analysis.

## Author Contributions

D. Mansour, R.C., A. Chandrasekaran, A.H.S., S.O.R., and P.R. designed the study. D. Mansour and A. Chandrasekaran designed the code and conducted simulations. D. Mansour prepared all the figures resulting from the simulations. R.C., T.M., D.K., A.K., and A. Chhabra conducted the biological experiments, while D. Milovanovic, S.K., A.H.S., and S.O.R. supervised. D. Mansour, R.C. and S.O.R. analyzed the data. D. Mansour, S.O.R, and P.R. wrote the initial draft of the manuscript. D. Milovanovic, S.O.R, P.R. were responsible for funding acquisition. All authors edited and contributed to the manuscript.

## Supplementary Tables

**Supplementary Table 1:** Table of parameters used in the LAMMPS model.

| Parameter | Value | Notes |
| --- | --- | --- |
| Simulation Box Size | $25\sigma \times 25\sigma \times 25\sigma$ | **Figure 3C-J:** A simulation box size of $50\sigma \times 50\sigma \times 50\sigma$ used in the $2 \times 2 \times 2$ stitched simulations. |
| Temperature | 1.0 | Assumes that the thermal energy of the system equals the depth of LJ particle interaction potential. |
| Time step size | $\Delta t = 0.0005\tau$ | $\tau = \sigma\sqrt{m/\epsilon}$ |
| Reduced Masses | $m_{\text{synapsin}} = 1.0m$<br>$m_{\text{actin}} = 0.567m$<br>$m_{\text{PEG}} = 0.108m$ | $(= 74097/74097 = 1.0)$<br>$(= 42000/74097 = 0.567)$<br>$(= 8000/74097 = 0.108)$ |
| Volume Fraction of Synapsin | $V_{\text{frac,Synapsin}} = [1\%, 2\%, 3\%, 4\%, 5\%]$ | Varied throughout |
| Number of Actin Filaments | $N_{\text{actin}} = [20, 30, 40, 50, 60]$ | **Supplementary Figures 5-6:**<br>$N_{\text{actin}} = [15, 40]$ |
| Volume fraction of PEG | $V_{\text{frac,PEG}} = 3\%$ | Kept constant throughout |
| **Non-bonded Interactions (Lennard-Jones Potential)** |  |  |
| Synapsin-Synapsin | $\epsilon_{\text{Syn-Syn}} = 2.0$<br>$\sigma_{\text{Syn-Syn}} = 1.0\sigma$<br>$\sigma_{\text{cutoff}} = 2.5\sigma$ | **Figure 1B-C:** $\epsilon_{\text{Syn-Syn}} = [1.0, 1.5, 2.0]$;<br>**Figure 5:** $\epsilon_{\text{Syn-Syn}}$ is alternated between 1.0 and 2.0;<br>**Supplementary Figures 5-6:**<br>$\epsilon_{\text{Syn-Syn}} = [0.5, 1.0, 1.5, 2.0, 2.5, 3.0, 4.0, 5.0, 7.5, 10.0]$ |
| Synapsin-Actin | $\epsilon_{\text{Syn-Act}} = 5.0$<br>$\sigma_{\text{Syn-Act}} = 1.0\sigma$<br>$\sigma_{\text{cutoff}} = 2.5\sigma$ | **Figure 5C-D:** $\epsilon_{\text{Syn-Act}} = 1.5$;<br>**Supplementary Figures 5-6:**<br>$\epsilon_{\text{Syn-Act}} = [0.5, 1.0, 1.5, 2.0, 2.5, 3.0, 4.0, 5.0, 7.5, 10.0]$ |
| Synapsin-PEG | $\epsilon_{\text{Syn-PEG}} = 1.0$<br>$\sigma_{\text{Syn-PEG}} = 0.9018\sigma$<br>$\sigma_{\text{cutoff}} = 1.0122\sigma$ | Purely repulsive interaction |
| Actin-Actin | $\epsilon_{\text{Act-Act}} = 1.0$<br>$\sigma_{\text{Act-Act}} = 1.0\sigma$<br>$\sigma_{\text{cutoff}} = 1.122\sigma$ | Purely repulsive interaction |
| Actin-PEG | $\epsilon_{\text{Act-PEG}} = 1.0$<br>$\sigma_{\text{Act-PEG}} = 0.9018\sigma$<br>$\sigma_{\text{cutoff}} = 1.0122\sigma$ | Purely repulsive interaction |
| PEG-PEG | $\epsilon_{\text{PEG-PEG}} = 1.0$<br>$\sigma_{\text{PEG-PEG}} = 0.8035\sigma$<br>$\sigma_{\text{cutoff}} = 0.9019\sigma$ | Purely repulsive interaction |

| Bonded Interactions (Harmonic Potential) |  |  |
| --- | --- | --- |
| Actin harmonic bond potential | Spring constant: 30,000 $k_B T/\sigma$<br>Equilibrium distance: $1.0\sigma$ | Determined empirically to ensure that the contour length of actin does not change within the droplet<sup>8</sup>. |
| Actin harmonic angle potential | Energy parameter: 2,500 $k_B T/\text{rad}$<br>Equilibrium angle: $180^\circ$ | Consistent with previous studies<sup>8,99</sup> |

**Supplementary Table 2:** Three-way ANOVA analysis of N_actin_, V_frac,Synapsin_, and the Order of Actin Addition.

| Source | SS | df | MS | F | P | $\eta_p^2$ |
| --- | --- | --- | --- | --- | --- | --- |
| $N_{\text{actin}}$ | 125.8620 | 4.0 | 31.4655 | 68.8051 | 2.0116e-45 | 0.3795 |
| $V_{\text{frac,Synapsin}}$ | 17.0451 | 4.0 | 4.2613 | 9.3181 | 3.0515e-07 | 0.07649 |
| Order of Actin Addition | 47.8796 | 1.0 | 47.8796 | 104.6974 | 3.1139e-22 | 0.1887 |
| $N_{\text{actin}} * V_{\text{frac,Synapsin}}$ | 9.0497 | 16.0 | 0.5656 | 1.2368 | 0.2357 | 0.04212 |
| $N_{\text{actin}} * \text{Order}$ | 10.5578 | 4.0 | 2.6395 | 5.7717 | 1.5475e-04 | 0.04880 |
| $V_{\text{frac,Synapsin}} * \text{Order}$ | 5.0138 | 4.0 | 1.2535 | 2.7409 | 0.02823 | 0.02378 |
| $N_{\text{actin}} * V_{\text{frac,Synapsin}} * \text{Order}$ | 9.0926 | 16.0 | 0.5683 | 1.2427 | 0.2314 | 0.04231 |
| Residual | 205.7912 | 450.0 | 0.4573 | - | - | - |

## Supplementary Figures

**Supplementary Figure 1:**
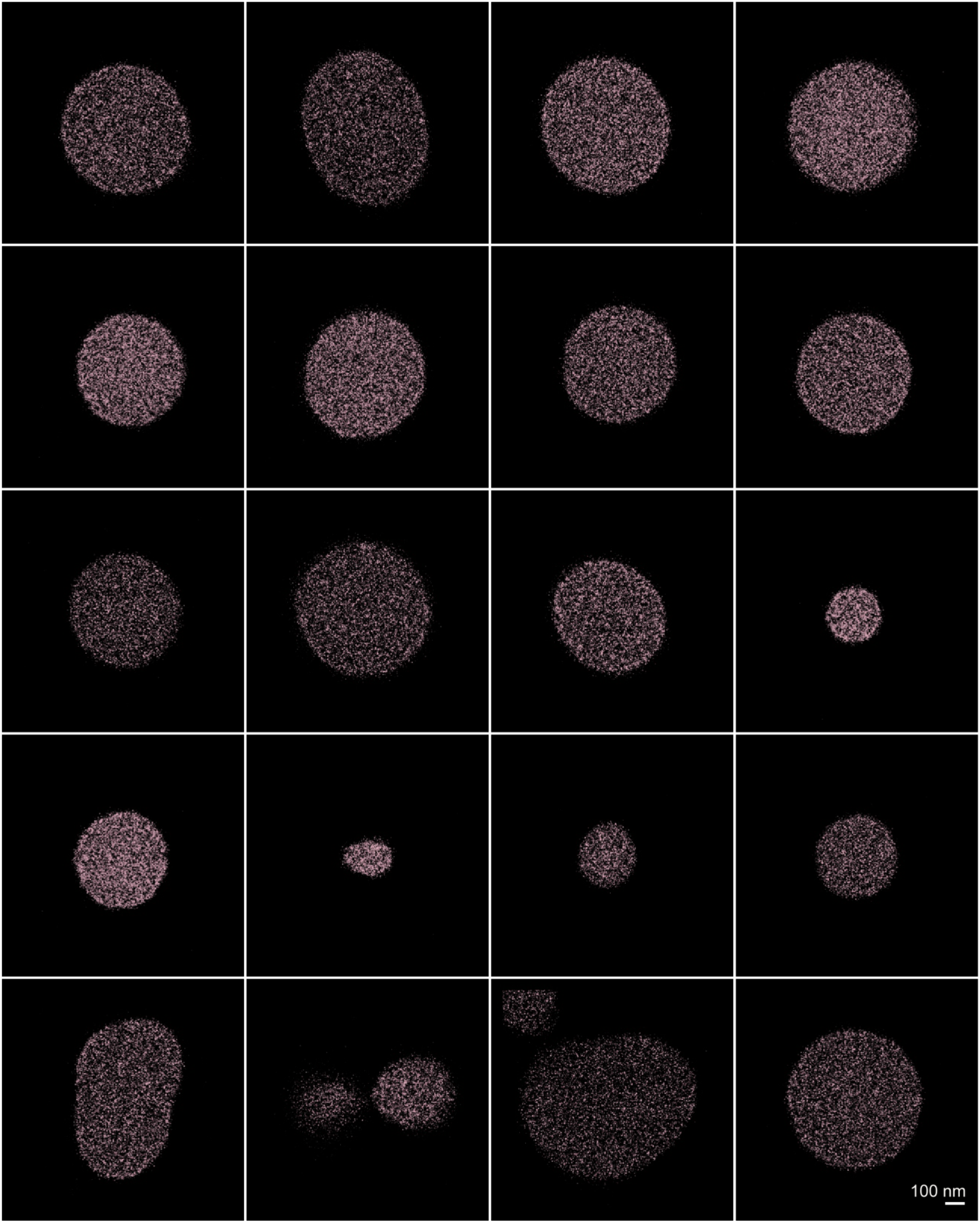
Image gallery of condensates consisting of synapsin alone (see **Fig. 4A** for details). Scale bar = 100 nm.

**Supplementary Figure 2:**
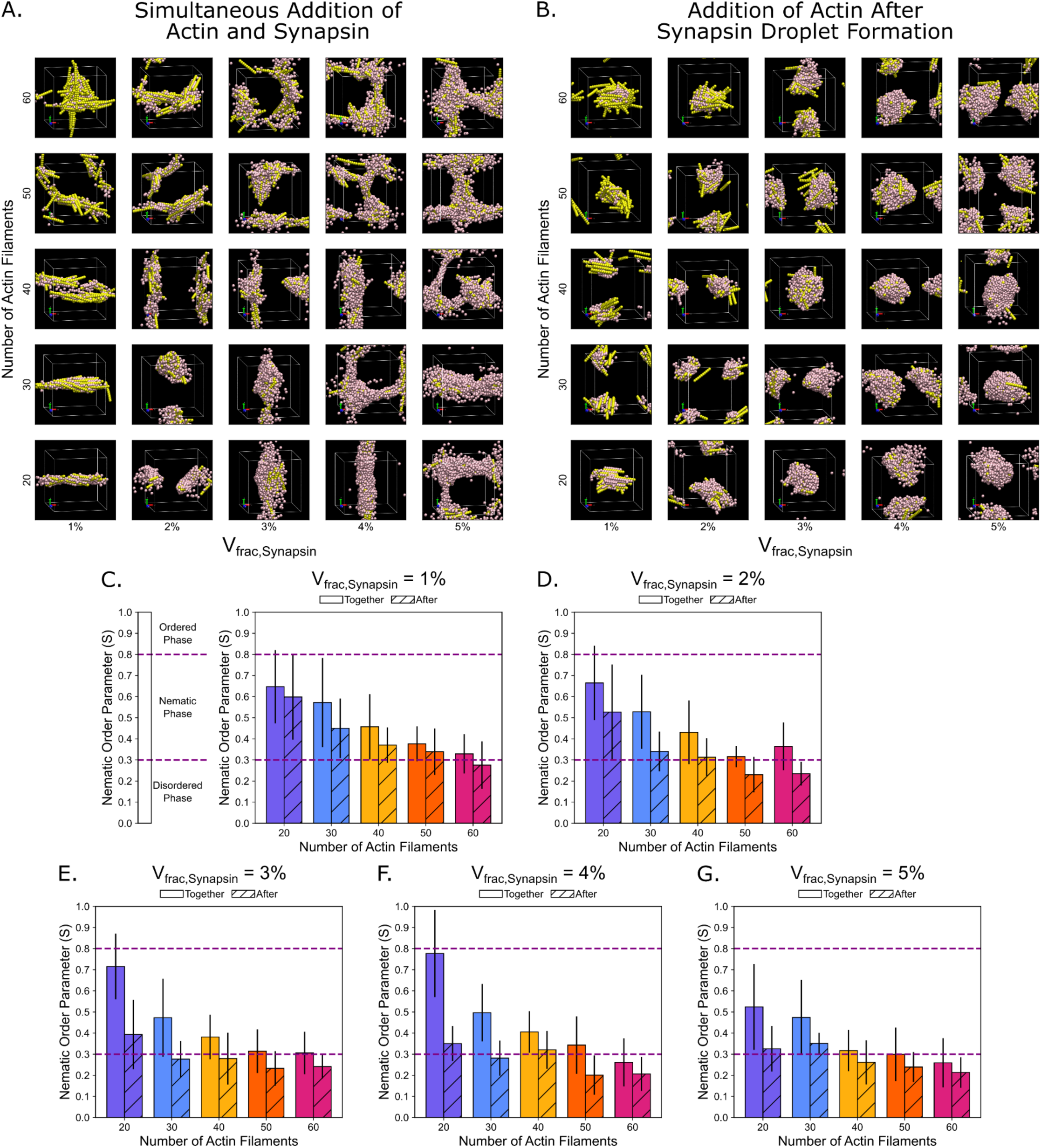
Actin-Synapsin composites result in the emergence of nematic order. **A)** Representative final snapshots for simulation conditions with the simultaneous addition of actin and synapsin. The number of actin filaments in the simulation is varied by row, while the volume fraction of synapsin is varied by column. The volume fraction of PEG is kept constant at 3%. PEG is not visualized in these snapshots. **B)** Representative final snapshots for conditions where actin is added to the system after synapsin droplet formation. The number of actin filaments in the simulation is varied by row, while the volume fraction of synapsin is varied by column. The volume fraction of PEG is kept constant at 3%. PEG is not visualized in these snapshots. **C-G)** Bar chart depicting the mean nematic order parameter and comparing between simulations where actin is added together with synapsin (empty bars) and after synapsin (diagonal stripes), for a synapsin volume fraction of **C)** 1%, **D)** 2%, **E)** 3%, **F)** 4%, and **G)** 5%. The error bars represent the standard deviation. *n* = 10 replicates, data from 10 time points in the last 100 frames of the simulations.

**Supplementary Figure 3:**
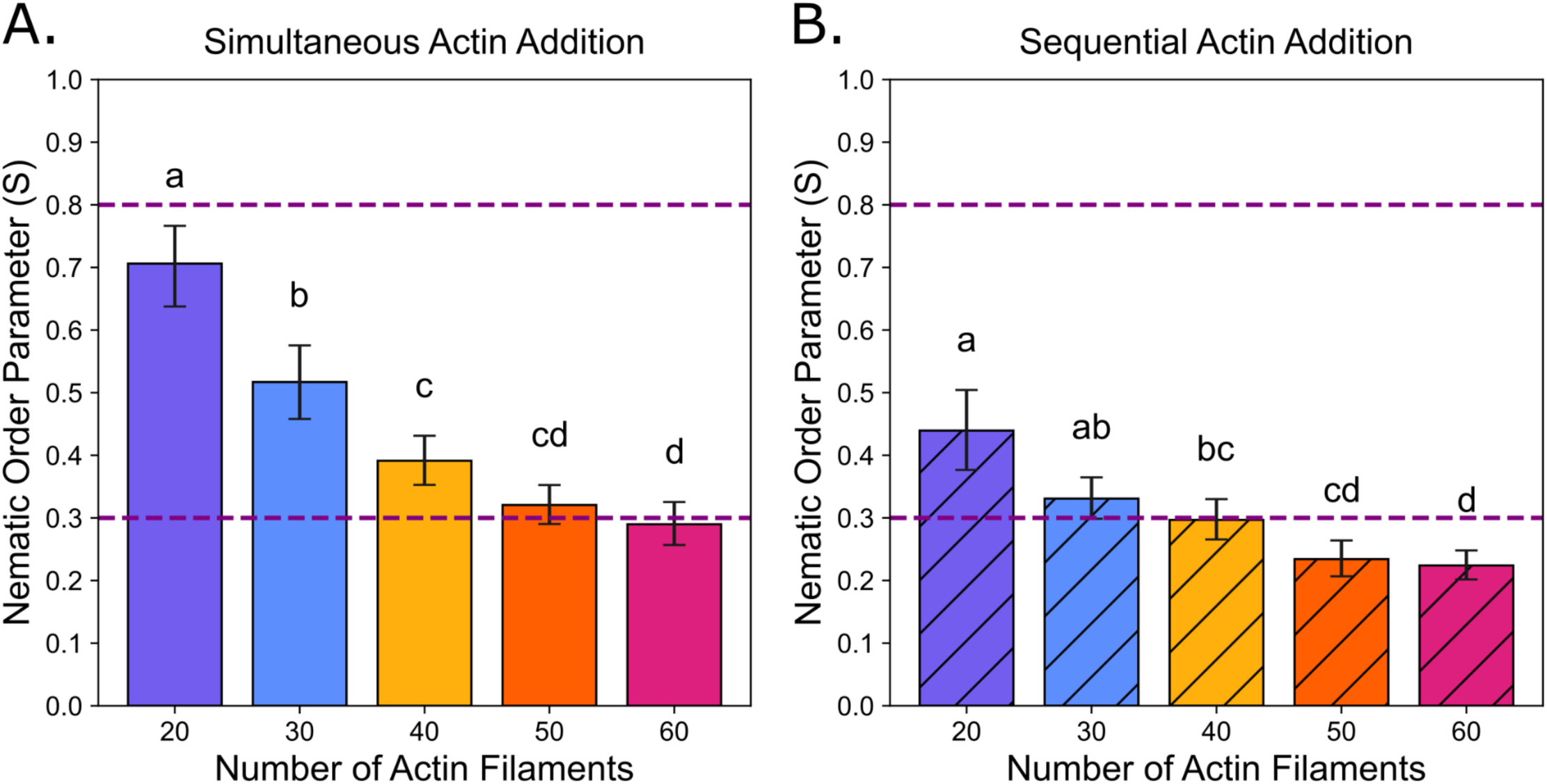
Average nematic order decreases as the number of actin filaments increases. **A-B)** Bar charts depicting the mean nematic order parameter for simulations where actin is added **A)** together with synapsin (empty bars) and **B)** after synapsin (diagonal stripes). Bar heights represent the mean nematic order reverse-transformed from the logit space. Error bars represent the reverse-transformed asymmetric 95% confidence intervals. Data for each N_actin_ condition is from data pooled across all V_frac,Synapsin_ conditions. Since each (N_actin_, V_frac,Synapsin_) condition has 10 replicates, each pooled N_actin_ condition has *n* = 50 data points. Statistical significance was evaluated using Games-Howell post-hoc tests conducted on the logit-transformed data. Please see **Supplementary Table 2** for the results of the three-way ANOVA. Distinct lowercase letters above each bar indicate the groups for the compact letter display. Groups with statistically significant differences (*P* < 0.05) do not share the same letters.

**Supplementary Figure 4:**
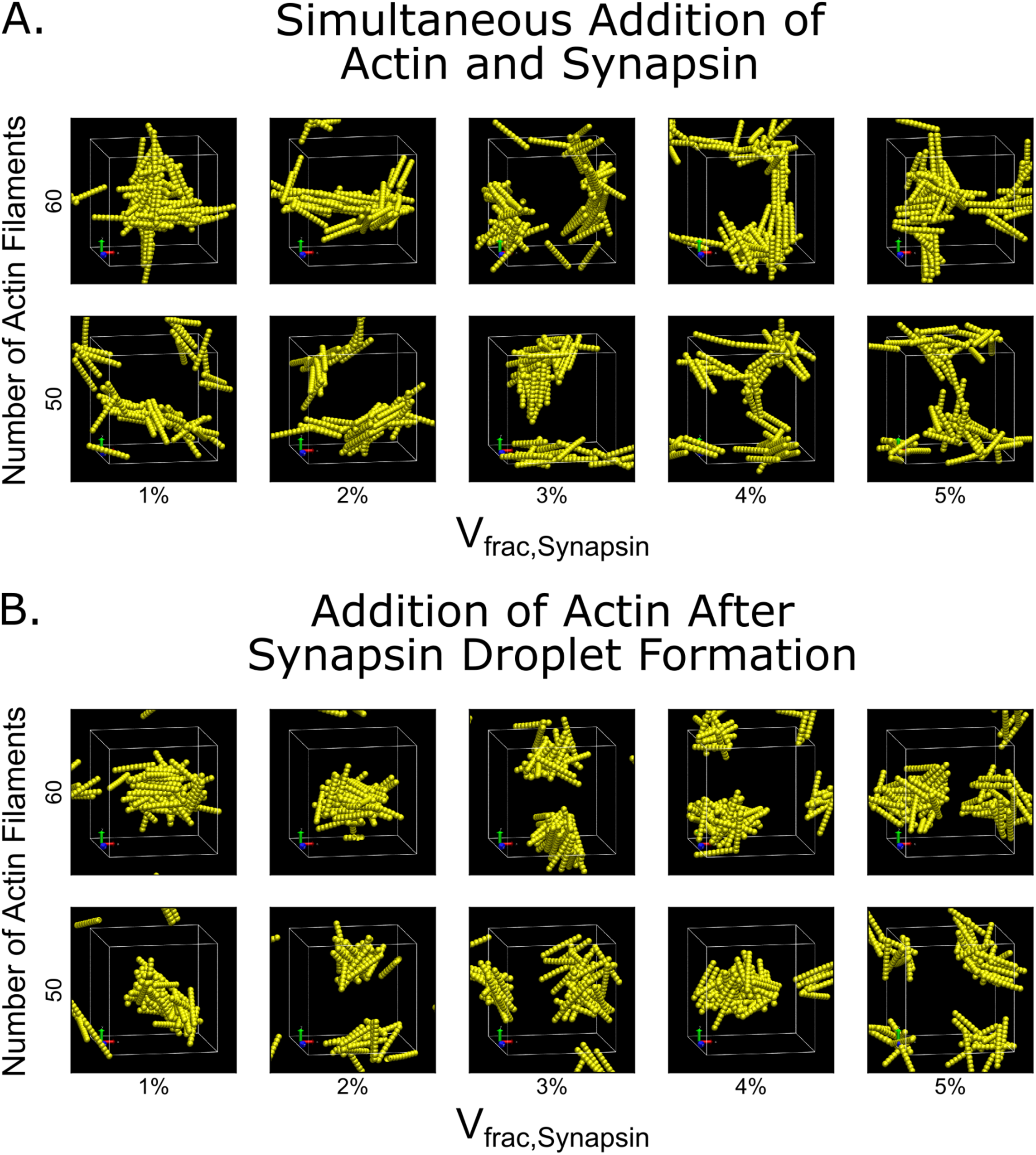
Gallery of images showcasing simulation conditions where actin filaments formed many intersections and became kinetically jammed. Representative final snapshots for simulation conditions with **A)** the simultaneous addition of actin and synapsin and **B)** where actin is added to the system after synapsin droplet formation. The number of actin filaments in the simulation is varied by row, while the volume fraction of synapsin is varied by column. The volume fraction of PEG is kept constant at 3%. Synapsin and PEG are not visualized in these snapshots.

**Supplementary Figure 5:**
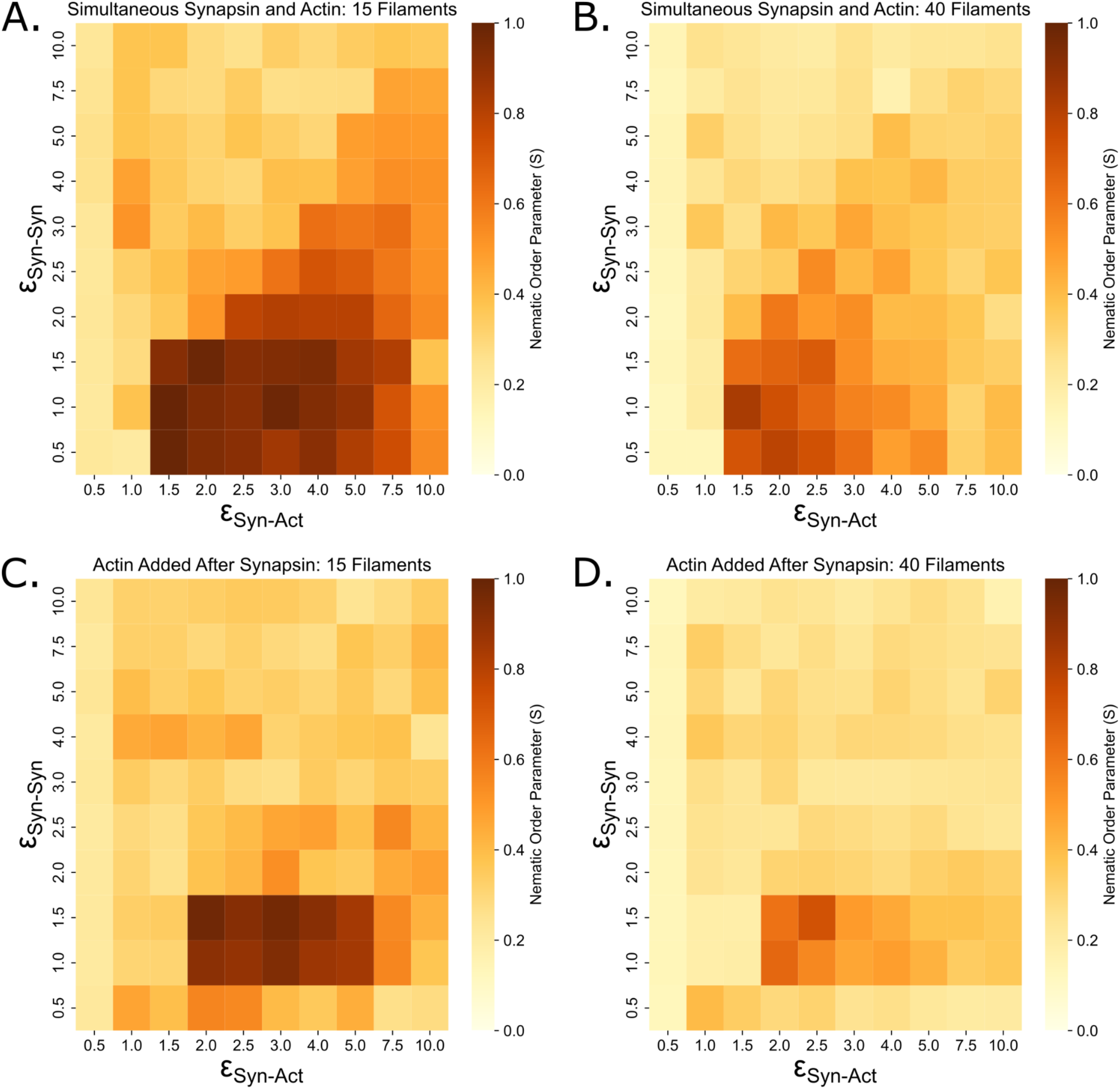
Energetic landscape of Synapsin-Synapsin and Synapsin-Actin interactions. **A)** Heatmap depicting the mean nematic order parameter for simulation conditions with 15 actin filaments added simultaneously with synapsin. **B)** Heatmap depicting the mean nematic order parameter for simulation conditions with 40 actin filaments added simultaneously with synapsin. **C)** Heatmap depicting the mean nematic order parameter for simulation conditions with 15 actin filaments added after synapsin droplet formation. **D)** Heatmap depicting the mean nematic order parameter for simulation conditions with 40 actin filaments added after synapsin droplet formation. For each condition, 3% synapsin volume fraction, 3% PEG volume fraction, and *n* = 10 replicates, data from 10 time points in the last 100 frames of the simulations. Please refer to **Supplementary Figure 6** for a representative gallery of final snapshots observed under the conditions studied.

**Supplementary Figure 6:**
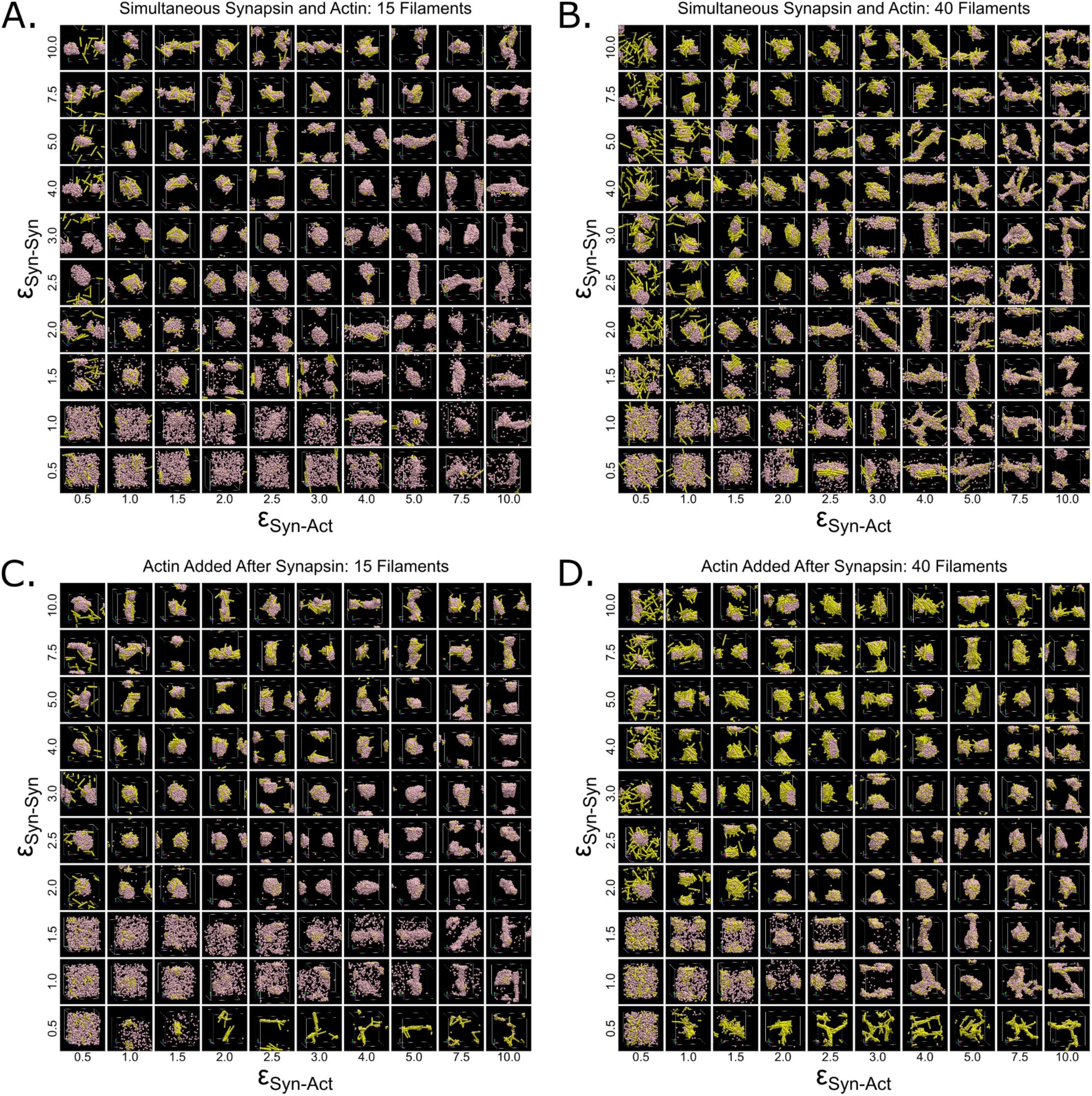
Simulation image galleries of energetic landscape of Synapsin-Synapsin and Synapsin-Actin interactions. Representative final snapshots for simulation conditions with **A)** 15 actin filaments added simultaneously with synapsin, **B)** 40 actin filaments added simultaneously with synapsin, **C)** 15 actin filaments added after synapsin droplet formation, and **D)** 40 actin filaments added after synapsin droplet formation. The number of actin filaments in the simulation is varied by row, while the volume fraction of synapsin is varied by column. For each condition, 3% synapsin volume fraction and 3% PEG volume fraction. PEG is not visualized in these snapshots. Please refer to **Supplementary Figure 5** for the nematic order analysis and heatmaps of the conditions studied.

**Supplementary Figure 7:**
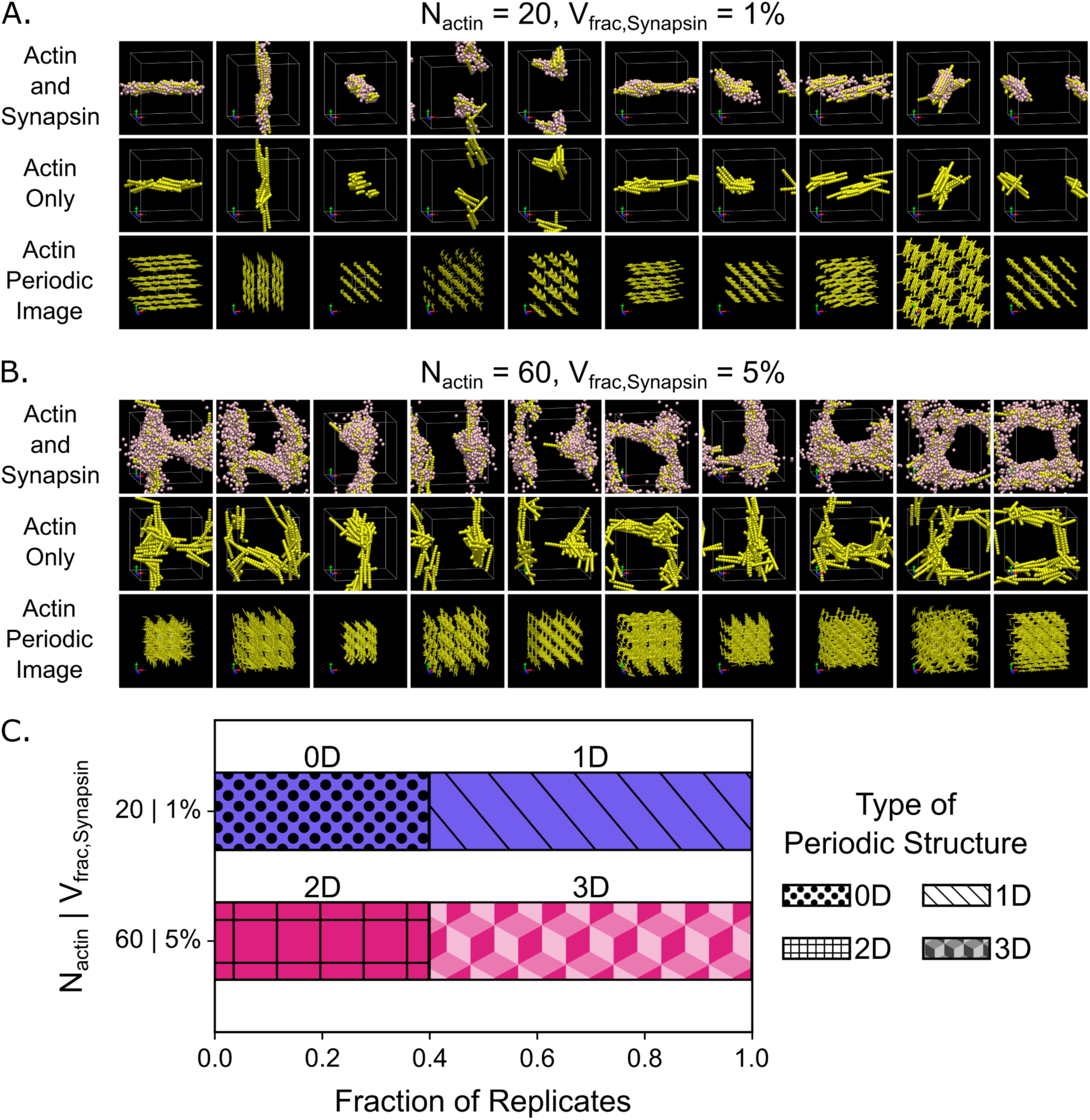
Comparison of actin structures formed in conditions that lead to high and low average nematic order. Gallery of final snapshots of all replicate simulations for conditions from **Figure 2A** and **Figure 3** with **A)** N_actin_ = 20 and V_frac,Synapsin_ = 1%, and **B)** N_actin_ = 60 and V_frac,Synapsin_ = 5%. Each column depicts the final snapshot of a single replicate simulation. The top row depicts the actin synapsin composite structure. The middle row depicts only the actin structure. The bottom row depicts the periodic images created by a 3×3×3 periodic extension of the simulation boxes to visualize the regularly organized structure formed by actin. PEG is not shown in any snapshots, and synapsin is shown only on the top row. **C)** Horizontal bar chart displaying a simple classification of the regularly organized structure formed by the replicates shown in panels **A** and **B**.

**Supplementary Figure 8:**
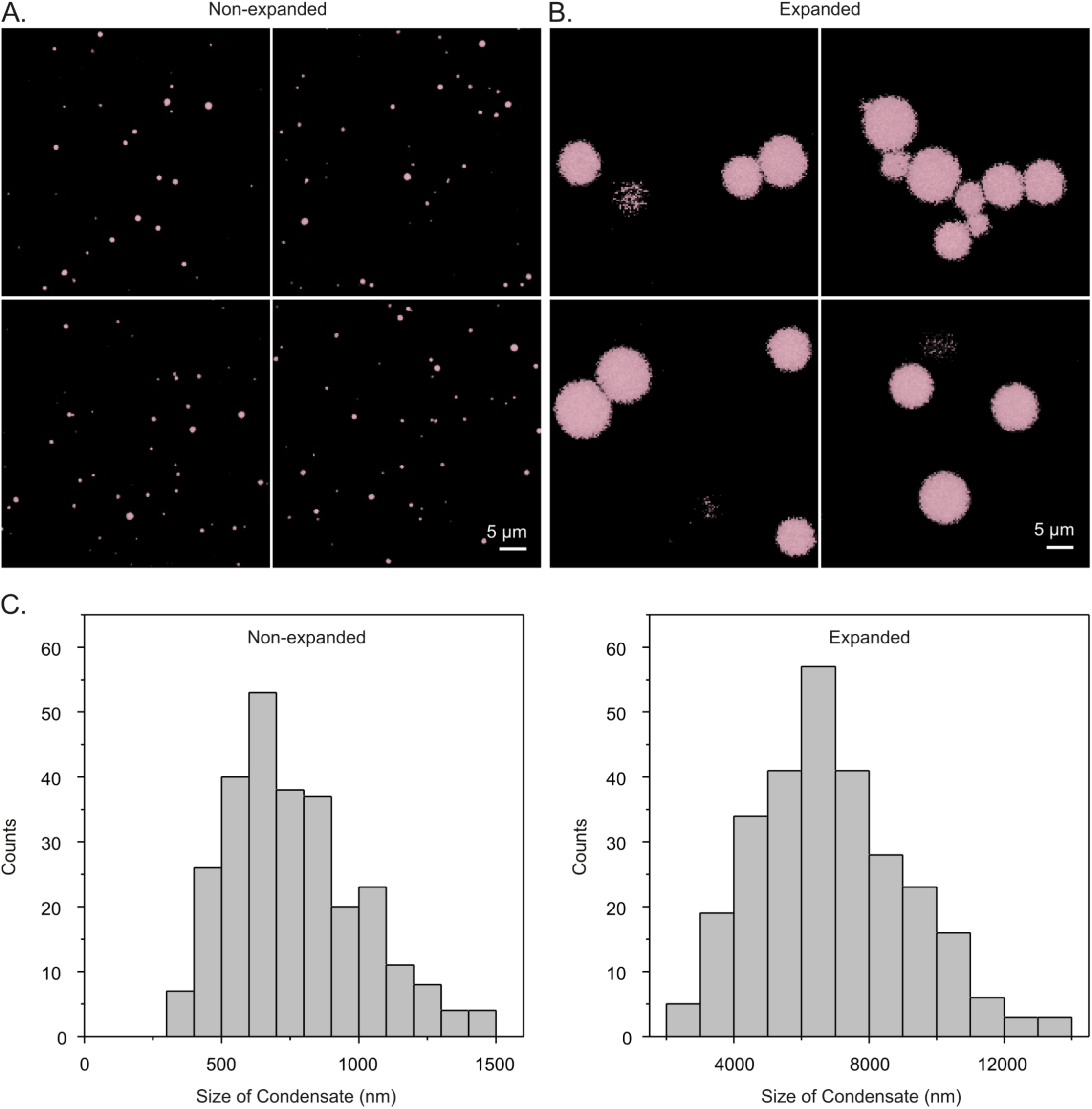
Expansion microscopy example for synapsin condensates. **A-B)** Non-expanded and expanded condensates are imaged using the same magnification. The size difference is obvious. Scale bars = 5 µm. **C)** Condensate size analysis. *n* = 271 codensates (non-expanded, average size = 724 nm). *n* = 276 condensates (expanded, average size = 6932 nm). The average expansion factor is 9.6 (based on *n* = 3 independent measurements).

**Supplementary Figure 9:**
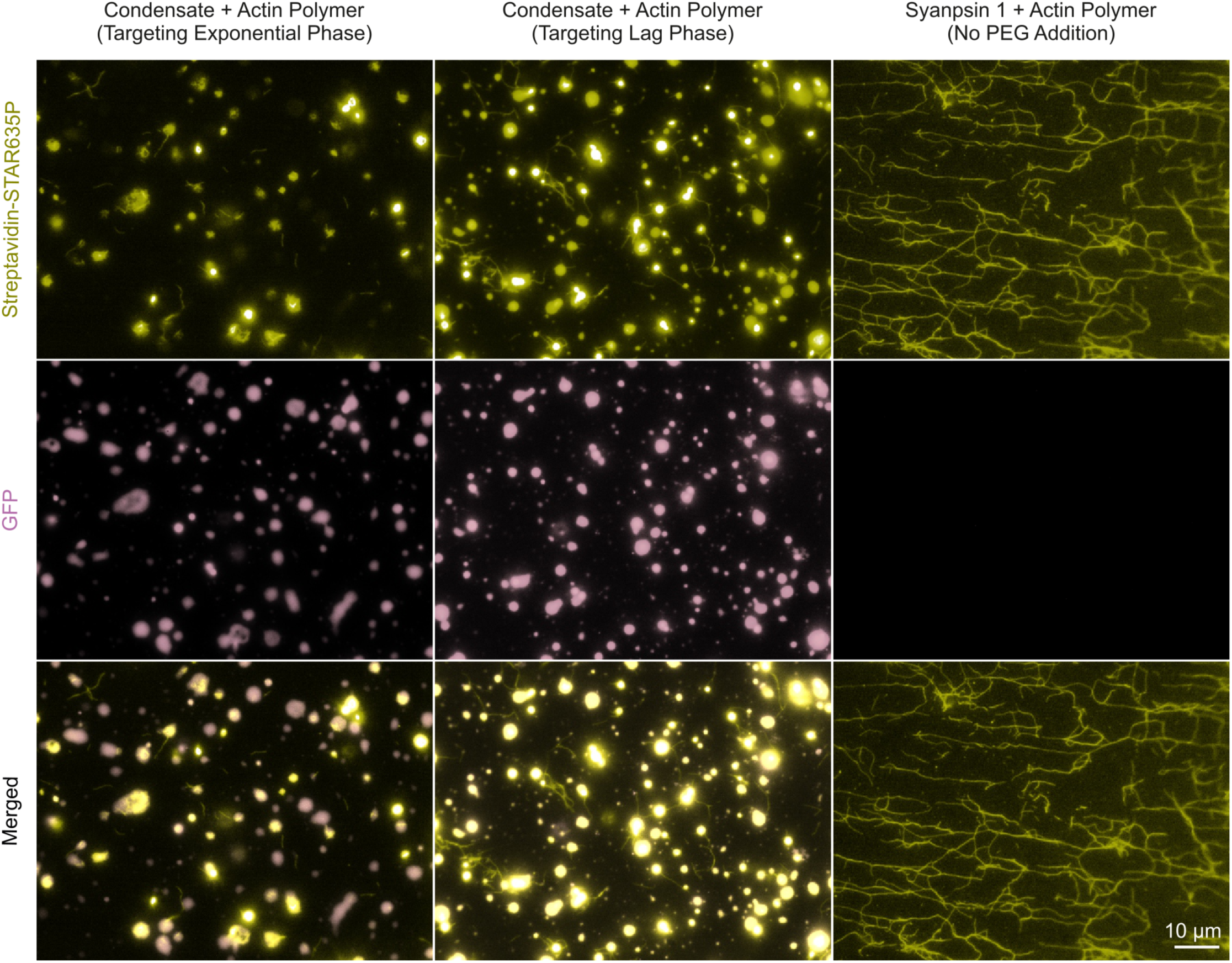
Conventional microscopy images of synapsin condensates containing actin filaments growing exponentially (left panels) or with a delay (middle panels). Incubating actin with synapsin, in absence of PEG for condensate induction, only results in the formation of actin filaments (right panels). Scale bar = 10 µm.

**Supplementary Figure 10:**
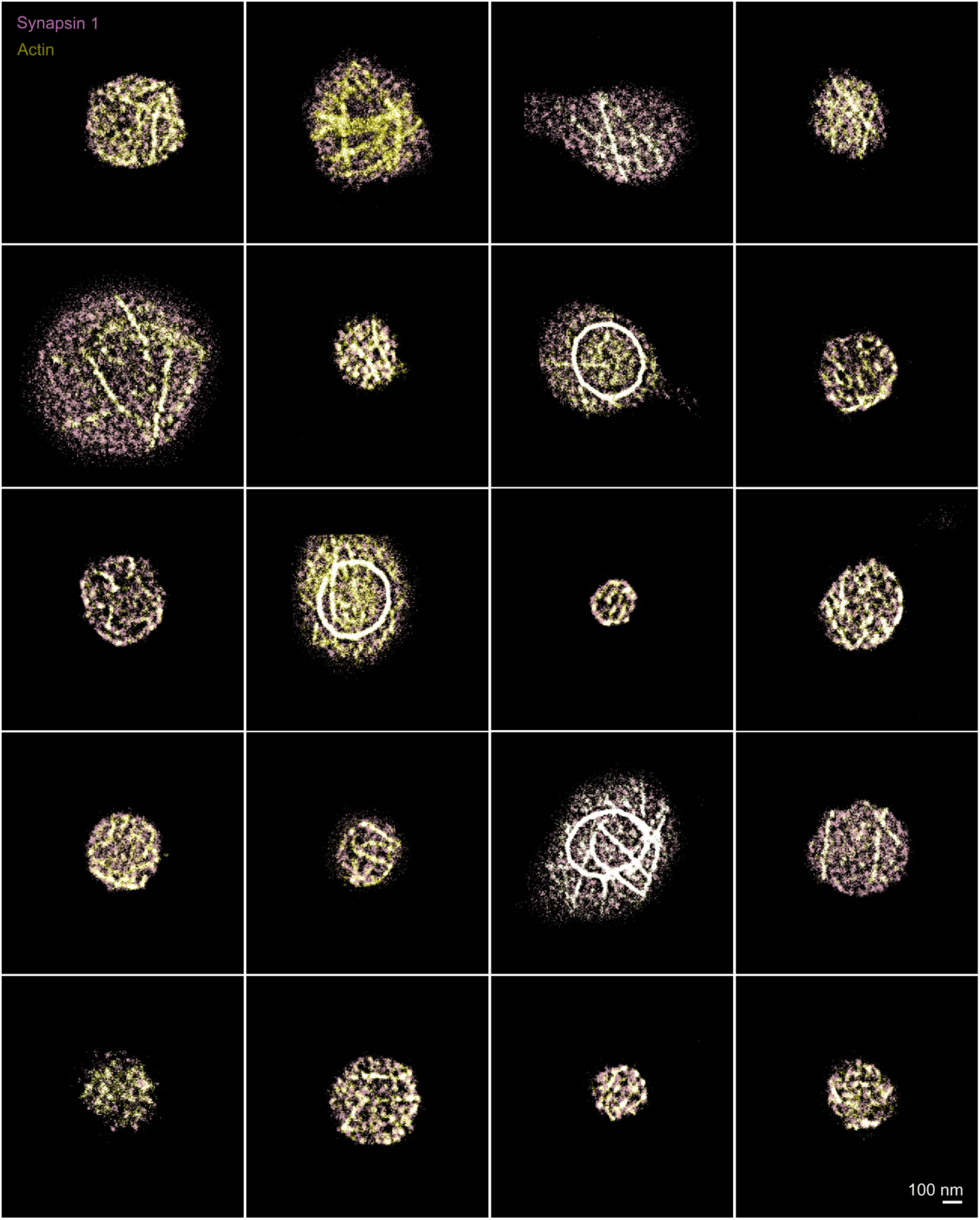
Image gallery of condensates consisting of synapsin and actin filaments growing exponentially (see **Figure 4B** for details). Scale bar = 100 nm.

**Supplementary Figure 11:**
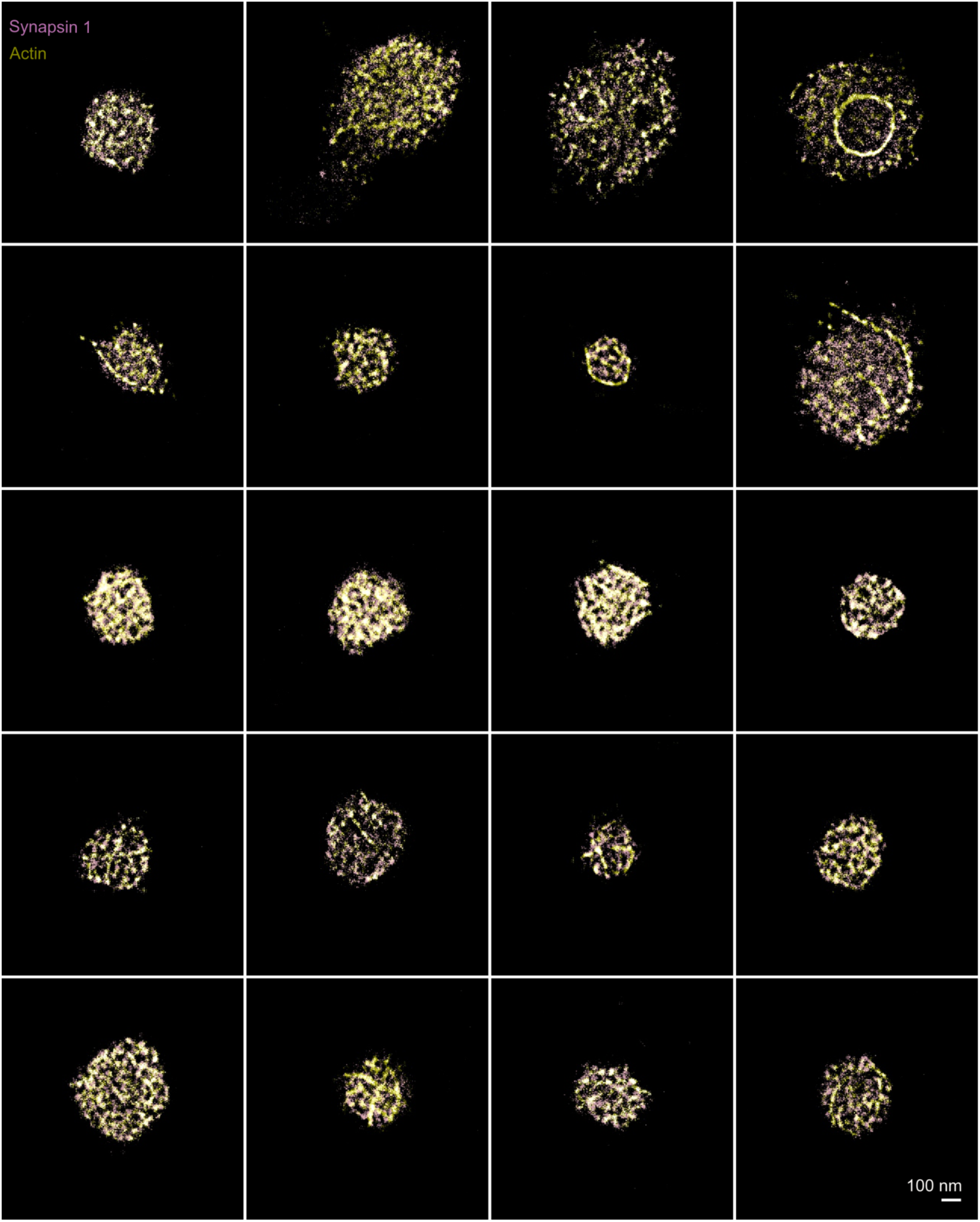
Image gallery of condensates consisting of synapsin and actin filaments growing with a delay (see **Figure 4C** for details). Scale bar = 100 nm.

**Supplementary Figure 12:**
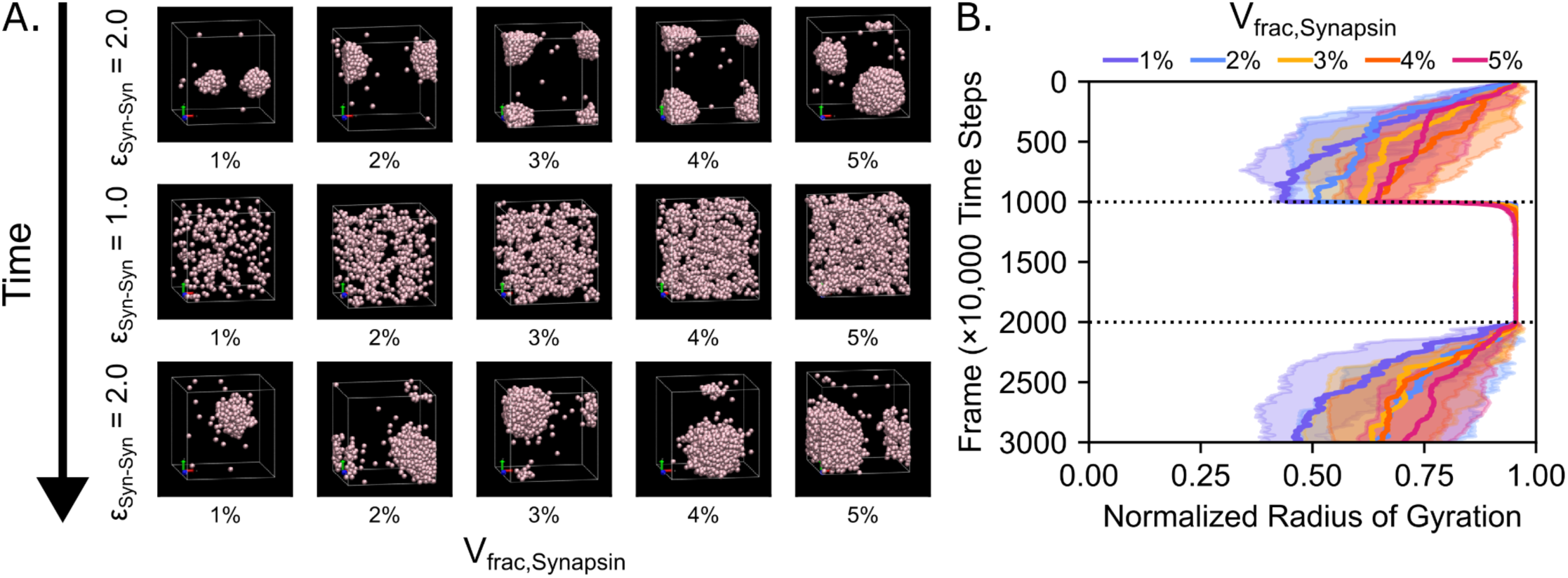
Altering the strength of Synapsin-Synapsin interactions can dissolve and reform synapsin droplets. **A)** Series of representative snapshots of simulations with synapsin and PEG particles depicting the initial formation, dissolution, and reformation of a synapsin droplet. ε_Syn-Syn_ is initially set to 2.0, lowered to 1.0 after frame 1000, and raised again to 2.0 after frame 2000. The volume fraction of synapsin is varied by column. The volume fraction of PEG is kept constant at 3%. PEG is not visualized in these snapshots. **B)** Time series showing the mean (solid line) and standard deviation (shaded area) of the normalized radius of gyration of the simulated synapsin particles, *n* = 10 replicates. The horizontal dotted lines indicate when ε_Syn-Syn_ was changed.

**Supplementary Figure 13:**
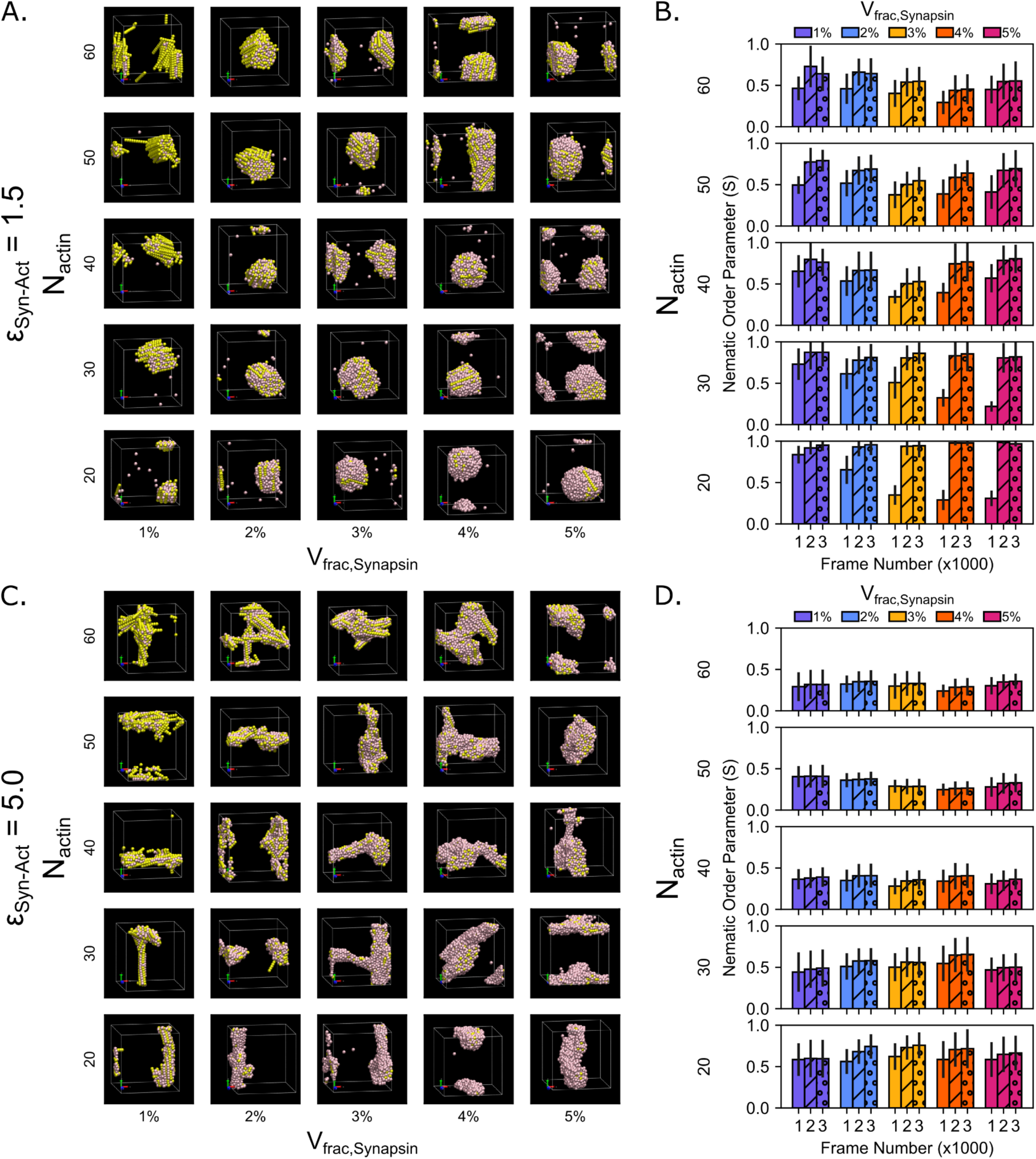
Many actin structures retain their organizational features following synapsin droplet dissolution. **A)** Representative final snapshots for simulation conditions with weak synapsin-actin attraction (ε_Syn-Act_ = 1.5). The number of actin filaments in the simulation is varied by row, while the volume fraction of synapsin is varied by column. The volume fraction of PEG is kept constant at 3%. PEG is not visualized in these snapshots. **B)** Bar chart depicting the mean nematic order parameter and comparing the simulation conditions in panel **A** at three illustrative time points (Frames 1000, 2000, and 3000) from the final frame of the three ε_Syn-Syn_ phases. Frame 1000 (empty bars) corresponds to the end of the first phase, where ε_Syn-Syn_ = 2.0. Frame 2000 (diagonal stripes) corresponds to the end of the second phase, where ε_Syn-Syn_ is decreased to 1.0. Frame 3000 (circles) corresponds to the end of the third phase, where ε_Syn-Syn_ is increased back to 2.0. The error bars represent the standard deviation. *n* = 10 replicates, data from 10 time points in the last 100 frames of the simulations. **C)** Representative final snapshots for simulation conditions with strong synapsin-actin attraction (ε_Syn-Act_ = 5.0). The number of actin filaments in the simulation is varied by row, while the volume fraction of synapsin is varied by column. The volume fraction of PEG is kept constant at 3%. PEG is not visualized in these snapshots. **D)** Bar chart depicting the mean nematic order parameter and comparing the simulation conditions in panel **C** at three illustrative time points (Frames 1000, 2000, and 3000) from the final frame of the three ε_Syn-Syn_ phases. Frame 1000 (empty bars) corresponds to the end of the first phase, where ε_Syn-Syn_ = 2.0. Frame 2000 (diagonal stripes) corresponds to the end of the second phase, where ε_Syn-Syn_ is decreased to 1.0. Frame 3000 (circles) corresponds to the end of the third phase, where ε_Syn-Syn_ is increased back to 2.0. The error bars represent the standard deviation. *n* = 10 replicates, data from 10 time points in the last 100 frames of the simulations.

